# The RRM domain-containing protein Rbp3 interacts with ribosomes and the 3′ ends of mRNAs encoding photosynthesis proteins

**DOI:** 10.1101/2024.07.09.602696

**Authors:** Luisa Hemm, Elisabeth Lichtenberg, Stefan Tholen, Viktoria Reimann, Kenta Kakazu, Sotaro Machida, Moontaha Mahbub, Oliver Schilling, Annegret Wilde, Satoru Watanabe, Conrad W. Mullineaux, Wolfgang R. Hess

**Author notes:** corresponding author: Wolfgang R. Hess **Email:**.

## Abstract

RNA recognition motif (RRM) domain proteins are crucial RNA-binding proteins (RBPs) across all domains of life. In cyanobacteria, single RRM domain proteins are involved in mRNA targeting to the thylakoid membrane and acclimation to certain stress conditions, but many details of their physiological functions and molecular targets have remained unknown. The model cyanobacterium *Synechocystis* sp. PCC 6803 has a family of three genes encoding the RRM domain-containing proteins Rbp1, Rbp2 and Rbp3. Here, we verified the RNA-binding activity of Rbp3 *in vivo* and show that cells of a Δ*rbp3* deletion strain had a lower PSI:PSII ratio and decreased pigment content and were significantly smaller than wild-type cells. To identify the set of interacting molecules, co-immunoprecipitation experiments were performed with a strain expressing a C-terminally FLAG-tagged Rbp3. Mass spectrometry of the elution fraction suggested physical proximity between Rbp3, ribosomes, and a very small number of other proteins. The most highly enriched transcript in the co-eluting RNA fraction was the *psaAB* mRNA. This was corroborated by fluorescent *in situ* hybridization (FISH) analyses showing decreased *psaA* mRNA signals in Δ*rbp3*, and colocalization with Rbp3-GFP in the wild type. Other mRNAs co-enriched with Rbp3 encode thylakoid, plasma membrane and carboxysome proteins. Binding assays using Bio-layer Interferometry validated the Rbp3-*psaAB* mRNA interaction, indicating a preference for folded RNA segments near or overlapping the respective stop codons.

**Significance statement:** The mechanisms by which proteins are produced at specific sites and inserted into the intricate membrane systems of photosynthetic cyanobacteria are only partially understood. While RRM domain proteins are well-studied RNA-binding proteins in eukaryotes, their functions in bacteria remain underexplored. This study reveals that the RRM domain protein Rbp3 in the cyanobacterium *Synechocystis* sp. PCC 6803 binds mRNAs encoding photosynthetic proteins, plasma membrane proteins and carboxysome proteins and localizes near ribosomes. The bound RNA segments are typically near the ends of coding regions, or in 5′ untranslated regions. These findings suggest that Rbp3 is involved in targeting mRNAs to various intracellular locations by interacting with structural elements within these mRNA molecules.

## Introduction

### RNA-binding proteins in bacteria

RNA-binding proteins (RBPs) play crucial roles in the physiological functions, processing and stability of RNA molecules. Although prokaryotes lack the extensive processing of pre-mRNAs that is typical of eukaryotes, they have a variety of RBPs that act in different steps of RNA-related processes. Among the best-studied RBPs in bacteria are the Rho factors, which terminate transcription (1), Hfq and ProQ, which mediate sRNA/mRNA interactions and can alter RNA stability (2, 3), and cold shock protein A, an important transcription antitermination factor (4). Little is known about putative homologs of these well-known RBPs in cyanobacteria. A structural homolog of Hfq exists in most cyanobacteria including the model *Synechocystis* sp. PCC 6803 (hereafter *Synechocystis* 6803), but the residues interacting with RNA in other bacteria are not conserved (5), and RNA-binding activity was not detected (6). However, systematic studies revealed several candidate RBPs in cyanobacteria (7–9). Some of these RBPs have recently been characterized, such as YlxR/RnpM interacting with RNase P (10), or PatR, a member of the larger OB-fold protein family binding several sRNAs and impacting the pattern of cell differentiation pattern in multicellular *Nostoc* sp. PCC 7120 (8), but only a few cyanobacterial RBPs have been analyzed in more detail thus far.

### RRM domain-containing proteins in cyanobacteria

The RNA recognition motif (RRM; also called ribonucleoprotein domain (RNP) or RNA-binding domain (RBD)) (11, 12) is the most common RNA-binding domain. It consists of a four-stranded antiparallel beta-sheet, stacked on two alpha helices (13). The domain was first described in proteins of heterogeneous nuclear ribonucleoprotein particles, where it is involved in the splicing of heterogeneous nuclear RNA (11). In addition to splicing, eukaryotic RRM proteins are now known to be involved in a variety of RNA-related processes. Eukaryotic RRM proteins are associated with mRNA stability (14), poly(A)-binding (15), mRNA transport (16), translation (17) and degradation (18). In prokaryotes, most proteins harboring an RRM domain are found in species in the phylum cyanobacteria (19). These bacterial RRM domain proteins typically have a single RRM domain, whereas eukaryotic proteins usually have multiple RRM domains.

RRM-domain proteins in cyanobacteria were discovered in *Synechococcus* sp. PCC 6301 (20), and homologs were soon found in many other cyanobacteria (21–24). Genes encoding these RBPs are strikingly conserved, with at least three members throughout the entire cyanobacterial phylum **(*SI Appendix*, Fig. S1)**, suggesting a function very relevant to the physiology of cyanobacteria. Distinct types of these RBPs have been recognized (25, 26), proteins with a C-terminal glycine-rich domain, frequently becoming expressed under cold stress (22, 27, 28), short proteins that consist of only the RRM domain and the longer Type II proteins with a relatively long C-terminal region of unknown function.

The presence of RRM domains in these proteins has prompted multiple attempts to verify their RNA binding experimentally. Sugita and Sugiura presented evidence of Rbp1 and Rbp2 from *Synechococcus elongatus* PCC 6301 to bind RNA *in vitro* (20). Utilizing a gel-filtration assay, RNA-binding activity was shown for the RBPs of *Anabaena variabilis* (27). Mutsuda et al. presented evidence of Rbp1 and Rbp2 binding to RNA *in vivo* based on native gel electrophoresis of nuclease-treated and -untreated cell extracts in *Synechococcus elongatus* PCC 7942 (29). Using an RNA-binding assay with homopolymeric RNA molecules, RNA-binding activity was demonstrated for RbpA1 of *Nostoc* sp. PCC 7120 *in vitro* (25). Mahbub et al. performed RT-PCR on RNA co-immunoprecipitated with Rbp2 and Rbp3 from *Synechocystis* 6803 (30), and Zhang et al. applied RNA-co-immunoprecipitation and high-throughput sequencing (RIP-Seq) to recombinant Rbp1 and Rbp2 from the same strain (26). Finally, evidence of *Synechococcus elongatus* PCC 7942 Rbp2 binding to RNA was obtained by incubating recombinant Rbp2 and poly(rU), followed by gel filtration (31). Based on these results gained for several different RBPs in various cyanobacteria, it is now timely to determine the population of interacting transcripts *in vivo*.

Three RRM domain-containing proteins are present in *Synechocystis* 6803, Rbp1, Rbp2, and Rbp3 (encoded by the genes *sll0517, ssr1480* and *slr0193*, respectively; ***SI Appendix*, Fig. S1**). Rbp1 is important under cold stress (32), although Rbp2 and Rbp3 are also somewhat cold-inducible (33). Rbp3 is involved in the acclimation to salt stress and the cold-induced lipid peroxidation (34). Indeed, the levels of some mRNAs encoding proteins involved in fatty acid biosynthesis and salt stress differed between a Δ*rbp3* mutant and the wild type, although the molecular basis of these differences remained unclear (34). Rbp3 consists of the RRM domain in the N-terminal 85 amino acids and an uncharacterized C-terminal section of 66 residues. It could be shown, that the salt tolerance in Δ*rbp3* mutants was restored when the Rbp3 C-terminus was fused to the N-terminus of Rbp1 suggesting that the salt-related aspects were not connected to the RRM domain (26). Conversely, the glycine-rich C-terminus of Rbp1 was dispensable under cold stress conditions (26).

Using fluorescent *in situ* hybridization (FISH) to probe the subcellular location of mRNAs encoding core subunits of the photosystems, we observed that Rbp2 and Rbp3 are involved in the targeting of *psaA* and *psbA* mRNAs towards the thylakoid membrane (30). Here, we analyzed the function of Rbp3 at the molecular and physiological level. We demonstrate actual RNA binding *in vivo* and show that this protein is relevant for the accumulation of a subset of mRNAs and determined the interacting proteins and transcripts. The data suggest that Rbp3 binds to the 3′ regions of mRNAs encoding photosynthesis core subunits and interacts closely with ribosomes. Our results further suggest specialized functions for the different members of this protein family, but point also at some functional complementarity. Finally, we present a working model for the function of Rbp3.

## Results

### Rbp3 mutant strains show differences in growth and pigment content compared to wild type

To phenotype the Δ*rbp3* deletion mutant, spot assays were performed in triplicate with stepwise 10-fold dilutions (**Fig. 1A, B**). Growth of the *Synechocystis* 6803 wild type on agar medium was compared to that of Δ*rbp3* and a complementation strain in which a plasmid pVZ322-localized *rbp3* copy was introduced into Δ*rbp3* and expressed from its native promoter. Growth of the Δ*rbp3* strain appeared slightly impaired compared to the other two strains. However, closer inspection revealed similar numbers of colonies in the three strains, while the colony sizes in Δ*rbp3* were significantly decreased compared to the wild type and complementation strain. After 8 days of growth under standard conditions, wild-type colonies had a median colony size of ∼0.6 mm, while that of the Δ*rbp3* mutant was ∼0.36 mm. This effect was rescued in the complementation strain Δ*rbp3+*Rbp3 under low light (**Fig. 1A**), and partially rescued under high light conditions (**Fig. 1B**). We conclude that the size, but not the number of colonies, was different in the absence of Rbp3 indicating that lack of Rbp3 slows growth, but does not induce cell death.

**Fig. 1.**
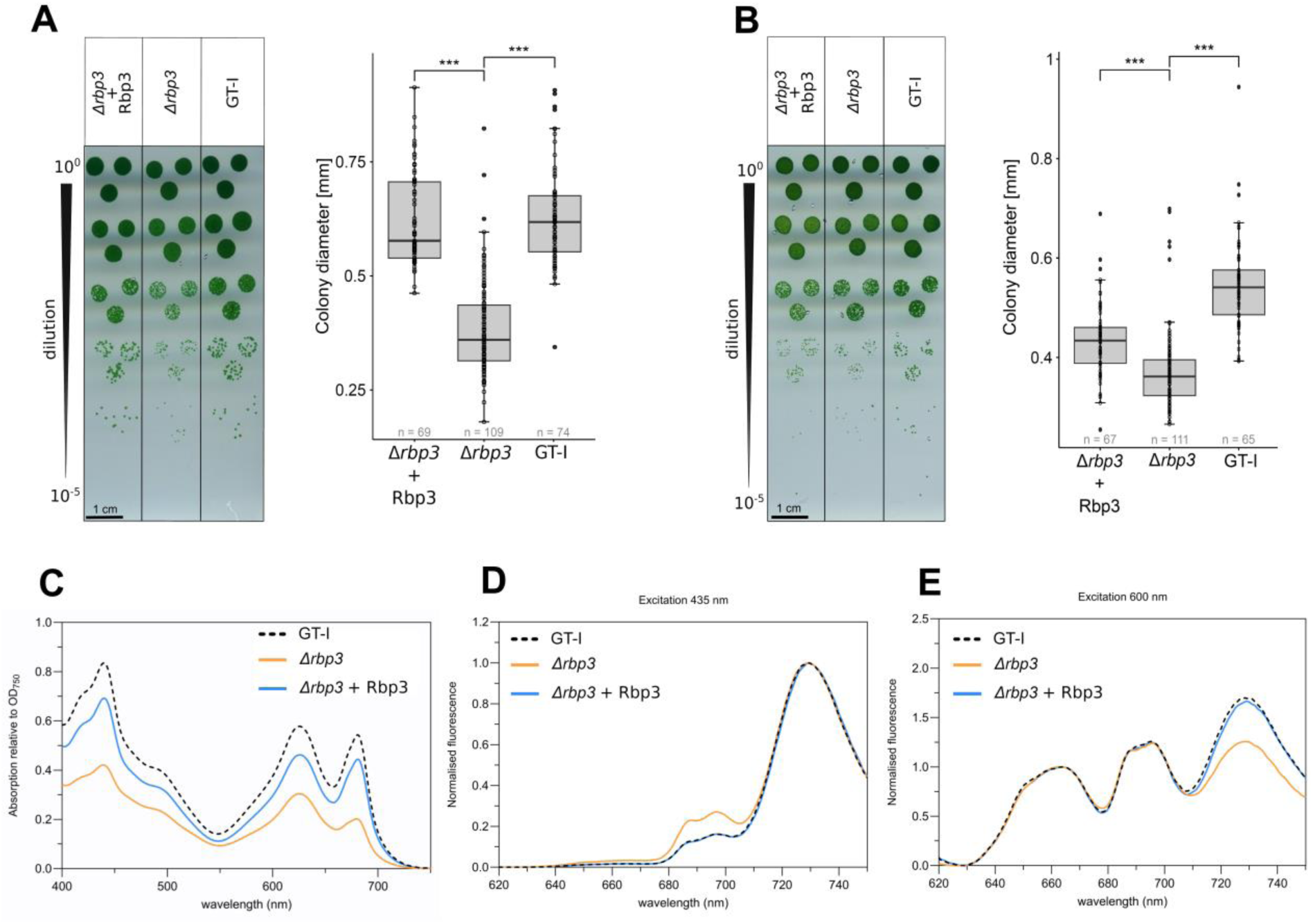
Phenotypical analysis of Rbp3 mutants. **(A)** Spot assays of the indicated *Synechocystis* 6803 strains after 8 days of incubation under standard growth conditions (40 µmol photons m^-2^ s^-1^, continuous light 30°C). **(B)** Spot assays under high light (300 µmol photons m^-2^ s^-1^) and 5 days of incubation. In both conditions, the growth of wild type *Synechocystis* GT-I strain was compared to the Δ*rbp3* deletion mutant and the complementation strain Δ*rbp3*+Rbp3 that expressed *rbp3* from an introduced pVZ322 plasmid under control of its native promoter. Culture aliquots were inoculated in triplicates and diluted ten-fold by every step as indicated. Colony sizes were measured and evaluated (right panels). Significances were calculated by the Students t-test, for the numeric values see ***SI Appendix,* Table S7**. **(C)** Room-temperature absorption spectra of cell suspensions for the GT-I wild type, Δ*rbp3* and Δ*rbp3*+*rbp3*, normalized at 750 nm. **(D)** 77K fluorescence emission spectra with chlorophyll excitation at 435 nm, normalized to the PSI peak. **(E)** 77K fluorescence emission spectra with phycocyanin excitation at 600 nm, normalized to phycobilin emission at 660 nm.

Whole-cell absorption spectra normalized to OD_750_ and 77K fluorescence emission spectra with chlorophyll excitation at 435 nm normalized to the PSI peak, or phycocyanin excitation at 600 nm indicated a lower cellular content of chlorophyll *a* and phycocyanin in Δ*rbp3* compared to the GT-I wild type (**Fig. 1C–E**). Moreover, these spectra indicated that the Δ*rbp3* cells had a lower PSI/PSII ratio. Introduction of an untagged Rbp3 restored spectra almost identical with the wild type (**Fig. 1C–E**). From these experiments, we conclude that the loss of Rbp3 affects the growth of cells as well as their photosynthetic properties, as indicated by the lower pigment content. This phenotype could be rescued by introducing a gene encoding Rbp3.

To allow the detection of Rbp3 by Western blotting and confocal microscopy, we complemented Δ*rbp3* mutants also with gene variants expressing a C-terminally fused 3xFLAG or sfGFP tag. To exclude that these tags altered the Rbp3 function, we verified complementation of the Δ*rbp3* mutant phenotype by expressing Rbp3-3xFLAG or Rbp3-GFP using absorption and 77 K fluorescence emission spectroscopy (**Fig. 2D, E** and ***SI Appendix*, Fig. S2).** Notably, the Δ*rbp3* cells were significantly smaller than the wild-type cells, a phenotype that was also complemented by the ectopic expression of Rbp3-sfGFP **(*SI Appendix*, Fig. S3).**

**Fig. 2.**
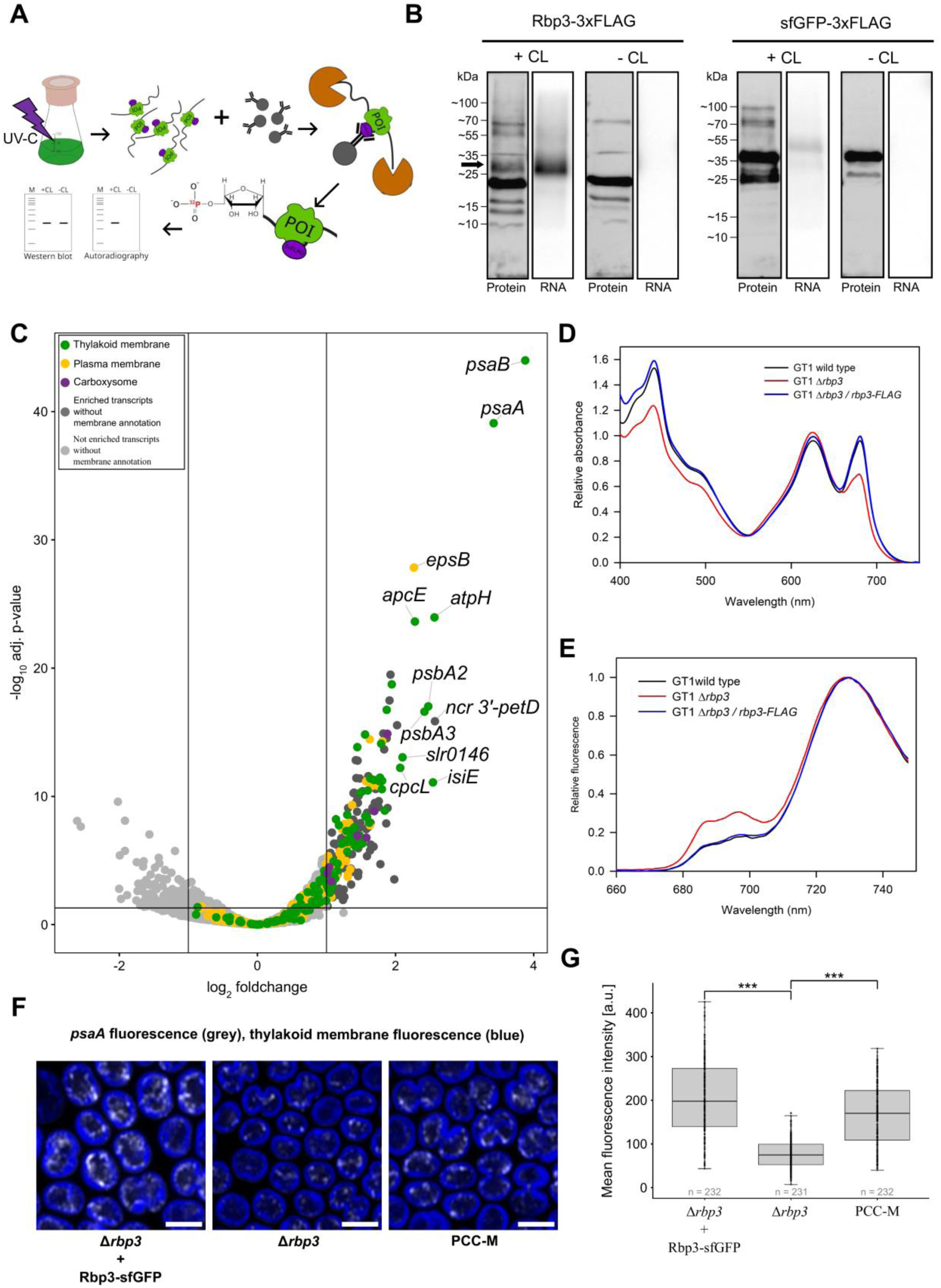
Verification of RNA-binding of Rbp3. **(A)** Schematic overview of the PNK assay (35). The protein of interest (POI) and RNA are crosslinked using UV-C. After cell lysis, co-IP is performed using anti-FLAG M2 magnetic beads. Non-covalently bound nucleic acids are removed by benzonase treatment. Nuclease-protected bound RNA is labeled with ^32^P using PNK. Purified proteins are separated by SDS-PAGE (78), transferred to a nitrocellulose membrane and visualized by autoradiography for ^32^P- labeled RNA and by western blotting for proteins. Controls omit the crosslinking step. **(B)** Rbp3 and sfGFP (negative control) PNK assays. Samples were cross-linked 24 h after induction of protein expression. After separating complexes by SDS-PAGE and western blotting, FLAG-tagged proteins were detected immunologically. Several bands appeared in the protein lanes (left subpanels). The most prominent band indicates monomeric Rbp3-3xFLAG at ∼20 kDa. For sfGFP-3xFLAG, the strongest signal is at ∼40 kDa, corresponding to monomeric sfGFP-3xFLAG. Bands below the signal of the respective monomer indicate degraded protein fragments, as these increased in intensity after UV treatment. Bands above the respective monomers belong to potential multimers, stabilized by UV treatment. In the UV-treated Rbp3-3xFLAG sample, a diffuse protein signal at ∼30 kDa (marked with an arrow) overlaps the autoradiography signal (left panel, + CL, RNA lane). The band at ∼70 kDa might belong to a multimer of Rbp3. **(C)** Volcano plot showing transcripts enriched in the Rbp3*_*3×FLAG CLIP RNA samples compared to the sfGFP*_*3×FLAG CLIP RNA samples. Colored dots indicate enriched transcripts encoding proteins involved in photosynthesis (green), proteins with the GO term membrane (yellow), associated with the carboxysome (purple), or others (dark gray). Light gray dots indicate non-enriched transcripts. Functionalities or gene names are given for transcripts enriched with a log_2_ fold change ≥2, the transcript for Rbp3 is not shown. (**D**) Room-temperature absorption spectra of cell suspensions for the GT-I wild type, Δ*rbp3* and Δ*rbp3*+*rbp3*-FLAG strains. **(E)** 77K fluorescence emission spectra, with chlorophyll excitation at 435 nm and normalized to PSI emission at 725 nm. **(F)** Confocal images showing the localization of *psaA* mRNA (TAMRA, grey) relative to the thylakoid membrane (chlorophyll, blue) in PCC- M (WT), *Δrbp3* and *Δrbp3* + Rbp3-sfGFP in the respective PCC-M WT background strain; scale bar = 2 µm (see ***SI Appendix,* Fig. S2** for the spectral characterization of the complementation strain expressing Rbp3-sfGFP and **Fig. S3** for the determination of cell sizes). **(G)** Quantification of *psaA* mRNA fluorescence intensity comparing PCC- M (WT), *Δrbp3* and *Δrbp3 +* Rbp3-sfGFP strains; *n* = number of cells. *** significant difference (*p* < 0.001). Significances were calculated by the students t-test assuming equal variances, for the numeric values see ***SI Appendix,* Table S8**.

### Verification of *in-vivo* RNA binding of Rbp3

Previous analyses suggested that Rbp3 is involved in the targeting of *psaA* and *psbA2* mRNAs towards the thylakoid membrane (30), but its actual RNA binding has not been directly demonstrated. To test the *in vivo* RNA binding of Rbp3, cultures expressing triple FLAG-tagged Rbp3 (strain Rbp3-3xFLAG) were used in an assay using the enzyme polynucleotide kinase (PNK) (35). The principle is outlined in **Fig. 2A**. After size-fractionating the complex and transfer to a nitrocellulose membrane, the labeled RNA was visualized by autoradiography and the protein by incubating the membrane with anti-FLAG antisera (**Fig. 2B**). For Rbp3, several bands were present in the protein panel, both for extracts from the crosslinked cultures and for non-crosslinked cultures used as negative controls. The most prominent band at ∼20 kDa corresponded to the molecular mass of the Rbp3-3xFLAG monomer (19.6 kDa). Below this, three other bands were observed, which were likely N-terminally truncated Rbp3-3xFLAG fragments. Two bands >20 kDa, one at ∼35 kDa and one at ∼70 kDa, were common to both Rbp3 protein lanes, that is, with and without UV treatment. For the crosslinked sample, two bands were specific: a somewhat diffuse band at ∼30 kDa and another at ∼55 kDa. In the autoradiogram, a single prominent band was observed at ∼30 kDa (**Fig. 2B**). Its position and somewhat diffuse appearance indicated that it belonged to the Rbp3-RNA ribonucleoprotein complex, which was also visible in the protein lane at the same molecular mass.

The sfGFP-3xFLAG control exhibited a prominent band just above the 35 kDa marker and several faint bands between ∼70 kDa and ∼100 kDa in the crosslinked protein sample, absent in the non-crosslinked sample, likely resulting from non-specific UV effects. A weaker band below the primary one indicated degraded sfGFP, implying UV treatment reduced protein stability. In the autoradiogram, only a very faint band was visible, possibly crosslinked sfGFP mRNA or RNA that had been near sfGFP during the crosslinking step. In comparison, the cross-linked Rbp3-3xFLAG sample showed a distinct band in the autoradiogram, matching a band in the western blot. This suggested that Rbp3 bound unknown RNA molecules *in vivo*, which could be isotope-labeled after protein purification and were protected from RNase digestion. The ∼10 kDa mass difference between RNA-free Rbp3 (∼20 kDa) and the RNA-Rbp3 complex indicated protected RNA fragments of approximately 60 nt. The diffused appearance of this band suggested a heterogeneous RNA population in length, composition, or both. The fact that Rbp3 was detected both with and without bound RNA suggested the existence of two pools of this protein.

### The majority of Rbp3 RNA interactors are transcripts encoding membrane-associated proteins

The results of the PNK assay suggested that Rbp3 was interacting *in vivo* with an RNA population heterogenous in length, composition, or both (**Fig. 2B**). To identify these transcripts, CLIP-Seq analyses were performed. Cultures expressing Rbp3-3xFLAG or sfGFP-3xFLAG were crosslinked using 1.25 J cm^-2^ UV-C. After enrichment of the samples with anti-FLAG magnetic beads **(*SI Appendix,* Fig. S4**), the protein was proteinase K digested and the RNA purified. On average, we obtained 1130 ± 269 ng of RNA from 200 mL of UV-treated Rbp3-3xFLAG cultures and 583 ± 107 ng from sfGFP-3xFLAG cultures. For control, we prepared total RNA from 25 mL aliquots of the same cultures after UV treatment. All samples were prepared in biological triplicates. After reverse transcription and sequencing, the RNA sequencing data were mapped to the genome of *Synechocystis* 6803, followed by a DESeq2 (36) analysis of transcripts enriched in the Rbp3-3xFLAG pulldown compared to the sfGFP-3xFLAG pulldown. A total of 229 transcripts were significantly enriched (log_2_FC > 1, *p* ≤ 0.05), including many mRNAs encoding membrane-associated proteins (**Fig. 2C**). Of these enriched transcripts, 41 were mRNAs encoding proteins associated with the thylakoid membrane, 21 encoded proteins associated with the plasma membrane, and seven proteins involved in carboxysome formation and carbon fixation. Additionally, 16 transcripts encode cytoplasmic proteins, for all other no location was annotated by a GO term. The transcripts with the highest enrichment were *psaA* and *psaB*, organized in a dicistronic operon and encoding the core proteins of photosystem (PS) I. Other highly enriched transcripts were *epsB* (*sll0923*) encoding a cell membrane protein, the tyrosine kinase Wzc related to the production of exopolysaccharides (37), *atpH (ssl2615)* encoding the small, membrane-intrinsic ATP synthase subunit c, *apcE* encoding the core-membrane linker polypeptide ApcE (or L_CM_), which anchors the giant phycobilisome light-harvesting structure to the membrane (38), *cpcL/cpcG2* encoding an alternative linker protein that facilitates energy transfer from a small phycobilisome type directly to PSI (39), and *psbA* encoding the PSII core protein D1. Additionally, the transcript for the cysteine-rich small protein IsiE, which is encoded in the *isiA*-*isiB* intergenic region (40) was highly enriched. For the complete list of 229 transcripts enriched in the co-IP with Rbp3 after UV cross-linking, see ***SI Appendix***, **Table S1**. In order to test whether deletion of *rbp3* had an effect on the *psaA* mRNA content per cell we used *in situ* FISH analysis. The results showed a significantly reduced *psaA* mRNA content per cell in Δ*rbp3* compared to the wild type and the complementation control Δ*rbp3+*Rbp3-sfGFP (**Fig. 2F, G**).

Based on these analyses, we conclude that Rbp3 binds several different mRNAs, the most abundant of which are the *psaA* and *psaB* mRNAs. The interaction of Rbp3 with these mRNAs is relevant for their accumulation level per cell and this should influence the number of PSI units.

### Specific transcript regions are enriched in the Rbp3 interactome

The enrichment of certain mRNAs in the analysis shown in **Fig. 2C** gives no information whether certain regions in these transcripts were more highly enriched than others. However, coverage plots of selected mRNAs indicated a highly uneven distribution of reads retained from crosslinking to Rbp3. For the dicistronic *psaAB* transcript, a peak was observed close to the 3′ region and overlapping the stop codon. Enrichment in regions close to the mRNA 3′ end was also observed for *psbA2* (***SI Appendix,* Fig. S5A, B**). In contrast, we observed for *rne* encoding RNase E, a clear peak in the middle of its 583 nt long 5′UTR, beginning exactly at the previously defined U-rich site (41) (***SI Appendix,* Fig. S5C**). A peak within the 5’UTR was also observed for *desC* (***SI Appendix,* Fig. S5D, E**).

Since no sequence similarity was detected between the peaks, we calculated the secondary structures of selected peak regions and the surrounding segments using RNAfold (42). For the transcript segments enriched for *psaAB* and *psbA2*, stem-loop structures were predicted that included the respective stop codon and short stretches of unpaired nucleotides. These structures are located upstream of the Rho-independent terminators (**Fig. 3B, *SI Appendix,* Fig. S5B**). For the element at the end of the *psaB* mRNA, we tested these interactions by Bio-layer Interferometry (BLI) using recombinant Rbp3 and biotin-labeled 50 nt ribooligonucleotides representing the most-enriched region of the *psaB* mRNA. For comparison, we tested the 50 nt sequences immediately before or after the peak segment (**Fig. 3C)**. The binding height towards the sequence of the coverage peak containing the STOP codon and the predicted Rbp3 binding site was higher than the binding height towards neighboring sequences (**Fig. 3D)**. To gain further insight in the likely interacting RNA motifs, we used RRMScorer, a recently developed algorithm that predicts the most probable interacting nucleotide segments (13). RRMScorer identified a 5′-UUNGN-3′ motif (*psaAB*: UUAGG and UUCGG; *psbA2*: UUGGU; *desC*: UUCGA) in the three shown enriched transcript segments with confidence scores between −0.248 and − 0.374, placing them clearly in the likely-binder region (13). These motifs include or are directly adjacent to unpaired nucleotide(s). Previously characterized RRM-domain containing RBPs bind single-stranded as well as structured RNA segments (13). Therefore, these results are in line with previous reports on other RRM-containing RBPs.

**Fig. 3.**
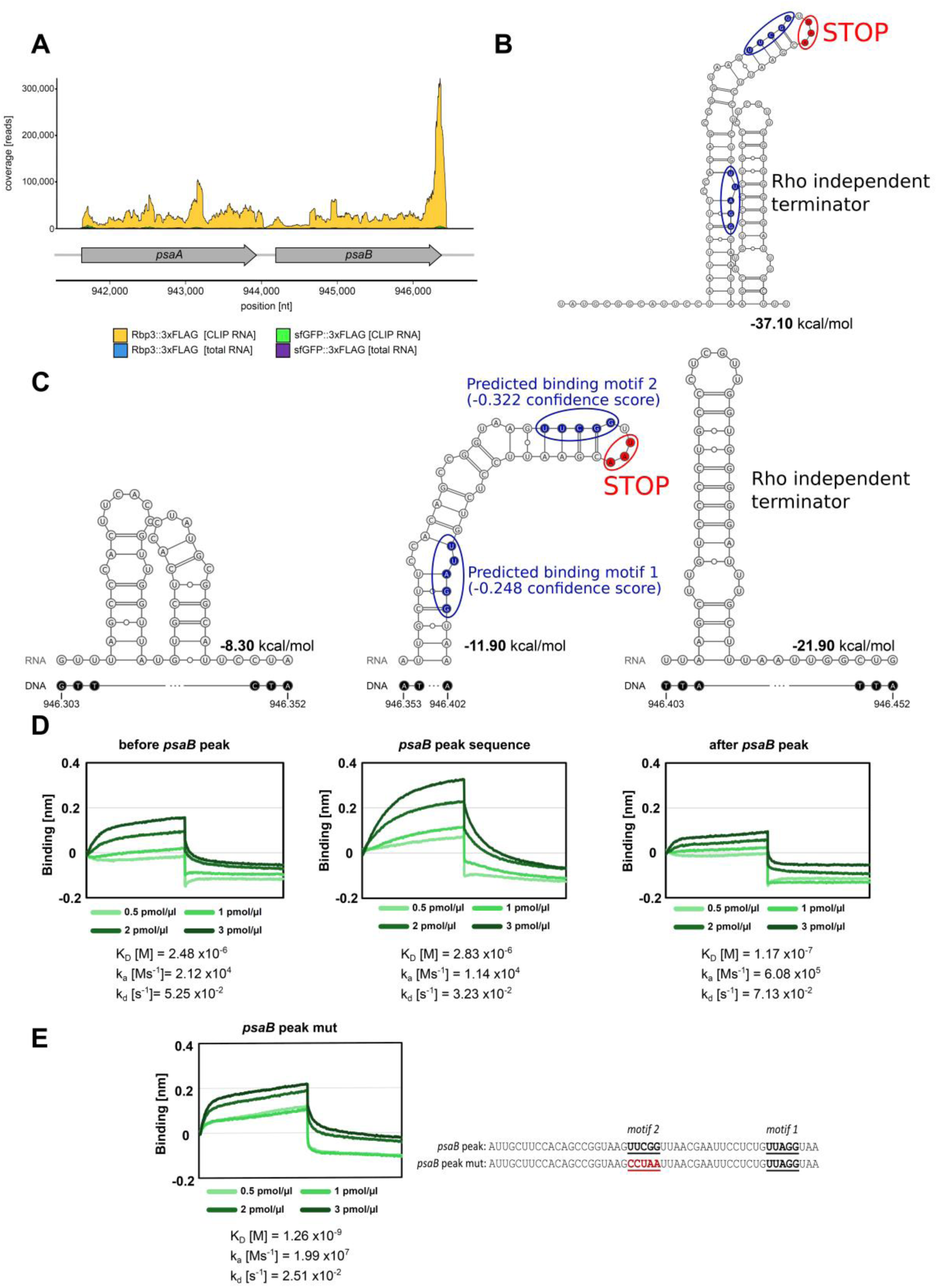
Coverage peaks in enriched mRNA reveal Rbp3 binding sites. **(A)** Read coverage along the *psaAB* locus in the Rbp3-3xFLAG CLIP RNA compared to the sfGFP-3xFLAG CLIP RNA (negative control), total RNA from cultures expressing the Rbp3-3xFLAG or sfGFP-3xFLAG. All cDNA libraries were sequenced in biological triplicates. **(B)** Secondary structure prediction around the *psaB* read coverage peak. **(C)** Three *psaB* mRNA regions were selected to test interactions with Rbp3. These regions encompass 50 nt of the *psaB* coverage peak (center), the preceding 50 nt of coding sequence (left), or the following 50 nt encompassing the Rho-independent terminator (right). (**D**) Recombinant Rbp3 was used in an *in vitro* Bio-Layer Interferometry assay to determine its binding affinities towards the 50 nt segments from panel C as synthetic RNA oligonucleotides. (**E**) Bio-Layer Interferometry assay with the *psaB* coverage peak RNA oligonucleotide in which the predicted binding motif 2 was mutated as shown. In panels B and C, the STOP codons are marked in red, and the putative RRM binding sites predicted by the RRMScorer algorithm are circled in blue (13). The secondary structures were calculated by RNAfold (42) and displayed using VARNA (95). For further mRNA coverage plots, see ***SI Appendix,* Fig. S5**.

The two high-scoring motifs predicted by RRMScorer for the *psaB* mRNA are located immediately before the stop codon (motif 2) and shortly after (motif 1; **Fig. 3B**), i.e. in a functionally sensitive region. To verify their relevance for the interaction with Rbp3, we mutated motif 2 (UUCGG) to CCUAA and performed another analysis by BLI. The result clearly showed a much-reduced interaction with Rbp3 (**Fig. 3E**). Therefore, motif 2 is indeed relevant for the interaction, either directly, or via an effect on the secondary structure, or both.

Different from these results, we observed enrichment along longer transcript stretches for mRNAs encoding carboxysome shell proteins or RuBisCO subunits (***SI Appendix,* Fig. S5F–H**).

### Rbp3 co-immunoprecipitates with the ribosome and LabA/NYN-domain proteins

The appearance of differently-sized signals for Rbp3 binding RNA and not binding RNA in our PNK assay (**Fig. 2B**) and the previous observation of involvement in thylakoid membrane targeting (30) posed the question if one or the other Rbp3 fraction would interact with other cellular structures. To identify possibly interacting proteins, co-immunoprecipitation (co-IP) analysis of Rbp3-3xFLAG was performed followed by mass spectrometry-based identification of the enriched proteins. A parallel analysis using sfGFP-3xFLAG as bait protein was used as control. Both analyses were done in triplicates and in total 686 proteins were detected. From these, 64 were significantly enriched (log_2_FC > 1, *p* = 0.05) for co-immunoprecipitating with Rbp3 (**Fig. 4, *SI Appendix,* Table S2**). Forty-one of these 62 proteins belong to the ribosome and were enriched with a log_2_FC > 2. Four ribosomal proteins could not or only be detected in one or two replicates in the sfGFP control (**Fig. 4B**). Two proteins, Slr1870 and Slr1859, were detected only in the Rbp3 pulldown. Slr1870 is an uncharacterized protein that harbors a LabA/NYN domain in the N-terminal and central two thirds, and an OST-HTH domain in the C-terminal third. We noticed that another highly enriched protein, Slr0650 (log_2_FC of +3.97), has a very similar structure. The enriched protein Slr1859 was characterized as an anti-sigma factor antagonist (43). Its gene *slr1859* belongs to a cluster that previously was characterized as involved in the regulation of carbon metabolism, particularly glucose metabolism (44). Besides the enrichment of ribosomal proteins, proteins involved in RNA degradation like RNase J (Rnj) or the polyribonucleotide nucleotidyltransferase (Pnp) were enriched. Another enriched protein was the flavodiiron protein Flv2, which is involved in alternative electron transport and photoprotection (45). The uncharacterized protein Sll0518 that harbors two oligonucleotide/oligosaccharide-binding (OB-fold) domains and is encoded directly downstream of Rbp1, had a log_2_FC of +3.69. In addition, seven more such potentially relevant other proteins were enriched, Slr0006, a TsaC fold protein involved in the N6- threonylcarbamoyladenosine modification of tRNA (46), the PilL-C protein Slr0322 (47), the thylakoid-associated ssDNA-binding protein Slr1034, as well as Slr0635, Slr1223 and Slr1854.

**Fig. 4.**
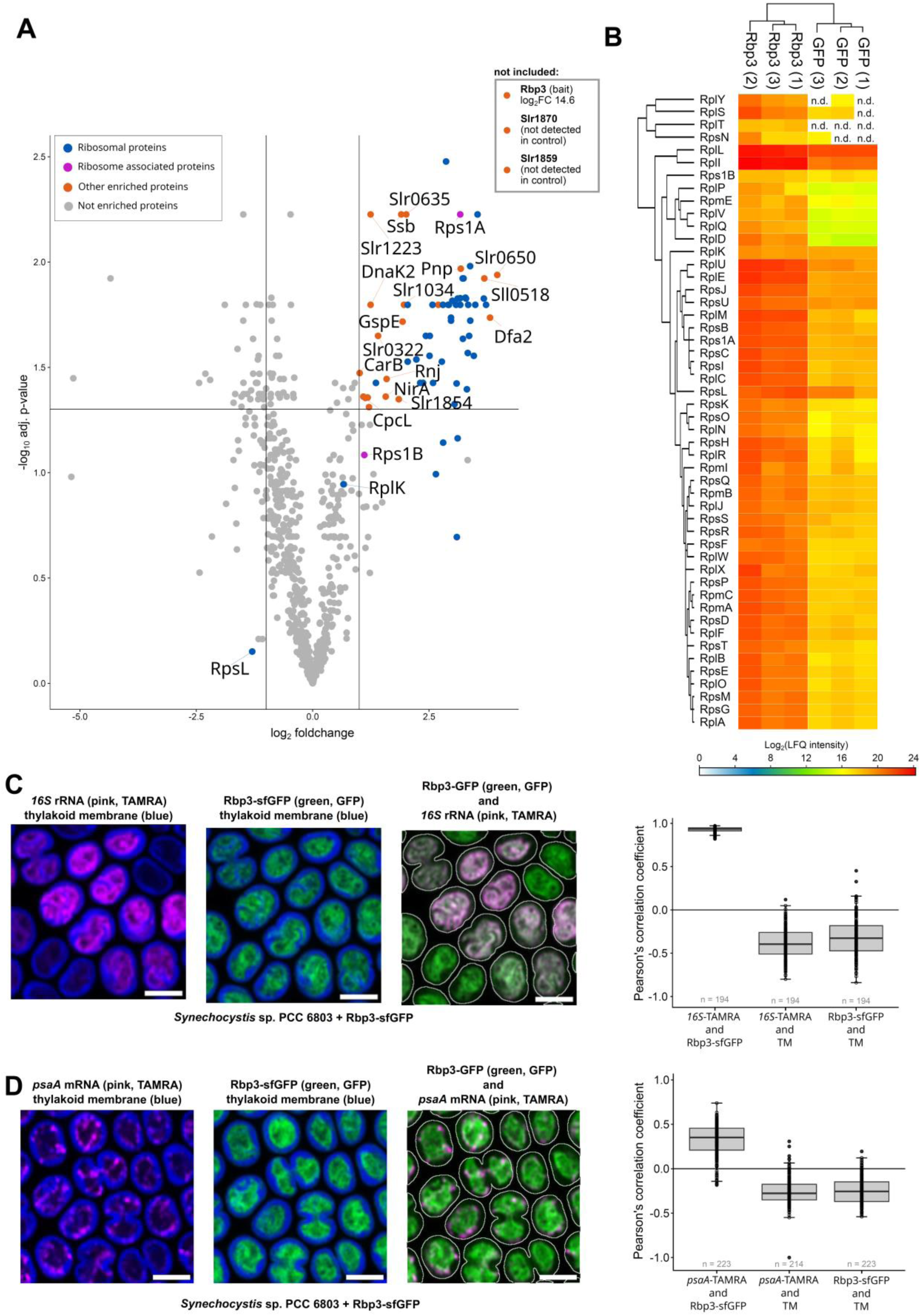
Protein interaction partners of Rbp3. **(A)** Volcano plot showing the protein interaction partners of Rbp3-3xFLAG identified by mass spectrometry compared to sfGFP-3xFLAG. Further details of the enriched proteins can be found in ***SI Appendix,* Table S2**. For the complete list of detected proteins, see ***SI Appendix,* Table S9**. **(B)** Heatmap showing the log_2_ (LFQ intensity) of all ribosomal proteins that could be detected. Hierarchical clustering was done using the Perseus software (96). Rbp3-3xFLAG and sfGFP-3xFLAG expression was induced 24 h prior to harvesting with 0.31 µM Cu_2_SO_4_. For the analysis, the Rbp3-3xFLAG as well as the sfGFP-3xFLAG bait protein were not included in the plot. Details of statistical analysis in ***SI Appendix,* Table S10**. **(C)** Left panel: Localization of hybridized 16S rRNA (TAMRA, magenta) and Rbp3-sfGFP (green) relative to the thylakoid membrane (blue) or the cellular outline (white) in *Δrbp3* + Rbp3-sfGFP (PCC-M background) cells. Right panel: Colocalization quantification of fluorescent pair (left panel, third image) by calculation of the Pearson’s correlation coefficient. *n* = number of cells. **(D)** Left panel: Localization of hybridized *psaA* mRNA (magenta) and Rbp3-sfGFP (sfGFP, green) relative to the thylakoid membrane (blue) or the cellular outline (white) in *Δrbp3* + Rbp3-sfGFP cells. Right panel: Colocalization quantification of fluorescent pair (left panel, third image) by calculation of the Pearson’s correlation coefficient. *n* = number of cells. Scale bar = 2 µm in panels C and D. For the numeric values in the calculation of statistical significances in panels C and D, see ***SI Appendix,* Table S11**.

The ribosome-associated protein Rps1A was significantly enriched, whereas Rps1B showed no significant enrichment due to its high abundance in the second replicate of the sfGFP pulldown. Both Rps1A and Rps1B were suggested to be involved in the Shine–Dalgarno-independent initiation of translation (48, 49).

To independently verify the association of ribosomes and Rbp3, RNA-FISH analysis of 16S rRNA was performed in cells expressing Rbp3-sfGFP. The Pearson’s correlation coefficient in the overlay between the sfGFP and the TAMRA (16S rRNA) signal was close to 1 (median: 0.935), strongly supporting the close interaction between ribosomes and Rbp3 (**Fig. 4C**). FISH labelling of the *psaA* mRNA in the same strain and microscopic analysis revealed a significant overlap with Rbp3-sfGFP fluorescence (**Fig. 4D**). The Pearson’s correlation coefficient in the overlay between the sfGFP signal and the *psaA* mRNA was positive (median: 0.352) supporting their interaction. That the correlation was not higher is consistent with the detection of two Rbp3 forms in the PNK assay, one binding RNA and the other non-binding (**Fig. 2B**), as well as the fact that our UV cross-linking experiment revealed interaction also with other transcripts than the *psaAB* mRNA (**Fig. 2C**).

### Transcriptomic analyses of the Δ*rbp2*Δ*rbp3* double mutant

To study the effect of Rbp3 deletion and to prevent complementation by Rbp2 we analyzed the transcriptome of the Δ*rbp2*Δ*rbp3* double mutant that was used previously in the investigation of mRNA thylakoid membrane targeting (30). The Rbps in cyanobacteria, especially Rbp1, are considered to be important factors in cold stress acclimation (19, 32). However, Rbp2 and Rbp3 are also somewhat cold-inducible (33, 34). Therefore, we analyzed the transcript levels 1 h, 2 h and 4 h after transferring the cultures from 30°C to 20°C. In addition, since the lack of Rbp3 affected photosynthetic properties (**Fig. 1C–E**), we performed an analysis 0.5 h, 1 h and 2 h following a shift to high light (250 µmol photons m^-2^ s^-1^). Subsequently, we performed microarray analyses of selected samples to identify the full set of transcriptomic differences.

### Transcriptomic differences in the comparison of Δ*rbp2*Δ*rbp3* versus wild type under non-stress conditions and after shift to 20°C

Effects on *psaAB* mRNA accumulation were already observed 1 h after cold stress induction and 0.5 h after shift to high light (**Fig. 5A, B**). Therefore, we focused on the comparison of transcriptomic data from these time points to the respective non-stress control. Under non-stress conditions (T0), the levels of 8 transcripts were significantly increased, and of 37 transcripts decreased in the *Δrbp2Δrbp3* to wild type comparison (**Fig. 5C, *SI Appendix,* Table S3**). The *psaA* and *psaB* mRNA levels (log_2_FC of −0.3) showed only a small decrease under these steady-state conditions (***SI Appendix,* Table S3**). The most decreased transcript levels (log_2_FC < 2.0), besides *rbp2* and *rbp3*, were observed for *sll0788*, which encodes a DUF305 membrane protein and the plasmid pSYSX-located genes *slr6040* and *slr6039* (*hik31a* operon), along with a slightly decreased level for the neighboring *slr6041* (**Fig. 5C**). The two *hik31* operons are involved in the control of heterotrophic growth, of acclimation to dark and low O_2_ conditions, and in the responses to several stress conditions (50). Moreover, the *hik31* operon is involved in copper resistance of *Synechocystis* 6803 (51). The eight upregulated transcripts included the complete transcriptional unit *pilA1*/*pilA2*/*sll1696* (***SI Appendix,* Table S3**). In addition, *ndhC* encoding the NDH-1 complex core subunit NdhC was upregulated (which was also enriched in the Rbp3 CLIP-Seq analysis, **Fig. 5C, *SI Appendix,* Table S1**). Interestingly, the *rbp1* transcript was upregulated when *rbp2* and *rbp3* were lacking, indicating a complementary function of the only remaining RRM-domain-containing protein. Furthermore, *slr0552* and *slr0601,* encoding proteins of unknown function and *slr0460* encoding an IS701-like transposase were upregulated.

**Fig. 5.**
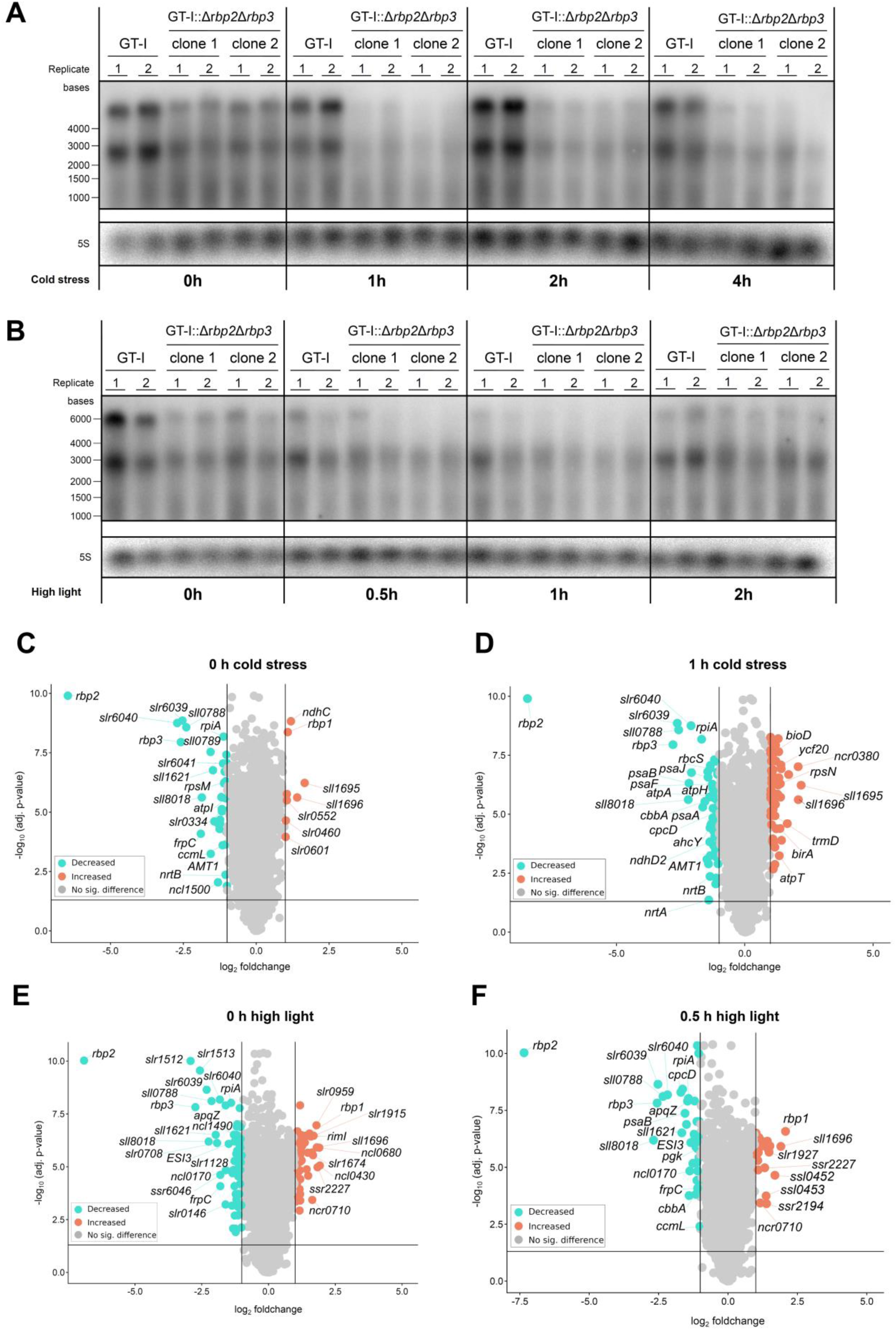
Transcriptomic analysis of Δ*rbp2*Δ*rbp3* double mutant. **(A)** Time course of *psaAB* mRNA accumulation in *Synechocystis* 6803 wild type (GT-I) or the Δ*rbp2*Δ*rbp3* mutant (clone 1, clone 2) after transferring the cultures from 30°C to 20°C for the indicated times. **(B)** Time course of *psaAB* mRNA accumulation after transferring the cultures from 50 to 250 µmol photons m^-2^ s^-1^. In both panels A and B, 5 µg of total RNA were separated on 1.5% formaldehyde-agarose gels and the RiboRuler High Range RNA ladder (Thermo Fisher) served as molecular mass standard. The single-stranded RNA probe encompassed the region from 100 to 252 nt after the *psaA* start codon. Hybridization against the 5S rRNA served as loading control. (**C, D, E and F)** Volcano plots showing transcripts with significantly increased or decreased levels in the comparison of the double mutant against wild type, 0.5 h after shift from 40 to 250 µmol photons m^-2^ s^-1^, or 1 h after shifting the cultures from 30°C to 20°C. The respective comparison at T = 0 h is also shown for both conditions. Transcriptomic changes were captured using a custom-made microarray. Salmon dots display increased transcript levels; turquoise dots display decreased transcript levels. Both experiments were performed with two replicates each. Note, that values for *rbp2* and *rbp3* represent background. They were included for completeness and do not indicate detected transcripts. See also **Tables S3** and **S4** and **Datasets 1** and **2** for the complete set of transcriptomic differences.

After 1 h of cold stress, 61 transcripts showed a significantly increased level and 64 a significantly decreased level in the double mutant compared to wild type (**Fig. 5D**). Notably, *psaA* (log_2_FC = −1.64) and *psaB* (log_2_FC = −2.17), the most enriched mRNAs in our CLIP-Seq analysis (**Fig. 2C**), showed the most decreased transcript levels. The levels of two other transcripts encoding PSI proteins were decreased in Δ*rbp2*Δ*rbp3* as well, *psaJ* (*sml0008*) with a log_2_FC of −1.43 and *psaF* (*sll0819*) with a log_2_FC of - 1.25. Other highly downregulated genes (log_2_FC < −2) were again the *hik31a* operon, two mRNAs encoding proteins of unknown function (*sll0788* and *sll8018*), as well as *sll1621* encoding peroxiredoxin. Several downregulated transcripts were identified as Rbp3 interaction partners in CLIP-Seq, such as *atpA*, *atpH, atpG* and *atpD* encoding ATP synthase subunits, mRNAs encoding components of the phycobilisome (*cpcD*), the cytochrome b_6_f complex (*petA* and *petD*), and proteins involved in carbon fixation (*rbcS* and *ccmM*) (***SI Appendix,* Table S3**).

Five transcripts showed highly increased levels (log_2_FC > 1.5, *p* = 0.05) in Δ*rbp2*Δ*rbp3* compared to wild type (**Fig. 5D**). Two of these transcripts (*pilA2* and *sll1696*) already showed higher levels under normal conditions. The other three transcripts included the sRNA *csiR1* and *slr1214* encoding the PatA-type response regulator NixC (47), also called LsiR (52). Gene *slr1214* is located downstream of the highly abundant *csiR1* sRNA and does not have an own promoter. The end of the *csiR1* transcript was significantly enriched in the Rbp3-3xFLAG CLIP-Seq pulldown, indicating a regulation at the RNA level (***SI Appendix,* Fig. S5I**). The full transcriptome dataset is visualized in **Dataset 1 Cold**.

### Transcriptomic differences in the comparison of Δ*rbp2*Δ*rbp3* versus wild type after shift to high light

The T0 samples from the cold shift and the light shift experiments were similar, but not identical due to different cell densities in the start cultures (OD_750_ of ∼ 1.0 in the temperature experiment and ∼0.6 in the light experiment). In cells grown under standard light conditions (T0), the steady-state levels of 58 transcripts were significantly increased and of 58 transcripts, besides *rbp2* and *rbp3,* significantly decreased (log_2_FC ≥ │1│, *p* = 0.05) in the double mutant compared to wild type (**Fig. 5E, *SI Appendix,* Table S4**). Notably, the *hik31a* operon, *sll0788, sll8018*, and *sbtA* and *sbtB*, encoding the bicarbonate transporter SbtA and its regulator SbtB (53), were strongly decreased.

Among the 58 transcripts with increased levels were three non-coding RNAs: *ncl0430*, a small asRNA transcribed from the complementary strand in the *psaA-psaB* intergenic spacer, *ncl0680,* constituting the 5′UTR of the *kaiC2B2* operon and *ncr0860* constituting the 5′UTR of the *hik34* gene (*slr1285).* Moreover, four transcripts encoding proteins of unknown function (*sll1696*, *slr0959*, *slr1915* and *slr1674*). Among the most increased transcripts were *slr0853* encoding the peptide-alanine-alpha-N- acetyltransferase RimI, *rbp1* and *ssr2227*, which encodes a putative transposase.

After 30 minutes of high light exposure (250 µmol photons m^-2^ s^-1^), the levels of 22 transcripts were significantly increased (log_2_FC > 1) and the levels of 40 transcripts decreased (log_2_FC < −1) (**Fig. 5F**). Again, the *hik31a* operon, as well as the transcripts of *sll8018*, *sll1621* and *sll0788* showed the strongest decreases. Additionally, transcript levels of the cell division gene *sepF* (54) were notably lower in Δ*rbp2*Δ*rbp3*. The transcript levels of some photosynthetic proteins showed a strong decrease (log_2_FC < −1.5), namely *cpcD*, *psaB* and *cpcC1*. These three transcripts were also significantly enriched in the Rbp3 CLIP-Seq dataset, as were other transcripts showing a decreased level upon light stress (***SI Appendix,* Table S1, S4**).

Of the 22 transcripts with increased levels, only four exhibited a log_2_FC > 1.5 and only *rbp1* had a log_2_FC > 2, underscoring the importance of the only remaining RRM-domain protein in Δ*rbp2*Δ*rbp3*. Among the transcripts with highly increased levels were furthermore the transcript for the phycobilisome degrading protein NblD (55) and the non-coding RNA *ncr0380*. The full transcriptome dataset is visualized in **Dataset 2 HL**. In summary, in cells deficient in both the *rbp3* and *rbp2* genes, the transcript levels of *psaA* and *psaB* were significantly lower under cold stress and high light stress compared to wild type (**Fig. 5**). The most pronounced reduction in *psaAB* transcript levels occurred 1 h after shifting from 30°C to 20°C (**Fig. 5A**). This decrease was detected by microarray analysis **(Fig. 5C–F, *SI Appendix,* Tables S3, S4**) and northern blot analysis (**Fig. 5A, B**). Transcript levels of several stress response proteins were also altered following *rbp2* and *rbp3* deletion.

## Discussion

All sequenced cyanobacterial genomes harbor a gene family encoding RRM domain-containing RBPs, typically consisting of three members, but sometimes more, such as eight in the N_2_-fixing, multicellular *A. variabilis* (27). RNA binding was previously verified for these eight Rbps using an *in vitro* assay (27). More recently, *in vitro* RIP-Seq analyses of *Synechocystis* 6803 recombinant Rbp1 and Rbp2 after total RNA incubation identified transcripts mainly related to iron uptake (26).

Here, we show that Rbp3, one of the three different RRM-type RBPs in *Synechocystis* 6803, binds RNA *in vivo* (**Fig. 2B**) and used CLIP-Seq to determine the interacting RNA population (**Fig. 2C**). The results are consistent with our previous report that Rbp3 is involved in targeting mRNAs encoding the core subunits of both photosystems to the thylakoid membrane, and demonstrate that Rbp3 physically interacts with them.

The lack of Rbp3 has consequences. Whole cell absorption spectra showed that the Δ*rbp3* mutant had a lower chlorophyll-to-phycocyanin ratio and a lower chlorophyll content per cell as suggested by the lower A_680_:OD_750_ ratio (**Fig. 1C, 2D** and **SI *Appendix*, Fig. S2A**). Furthermore, 77 K fluorescence emission spectra normalized to the PSI peak showed that the Δ*rbp3* mutant has a lower PSI:PSII ratio compared to the wild type (**Fig. 1D, 2E**and **SI *Appendix*, Fig. S2B**). Most of the chlorophyll in the cell is associated with PSI and therefore, this combination of effects is indicative of a reduced cellular PSI content (56, 57). This result is furthermore consistent with our finding that the *psaAB* mRNAs were the most-enriched transcripts in our CLIP-Seq analysis (**Fig. 2C**). Our CLIP-Seq results also identified several other mRNAs interacting with Rbp3. Most prominent were mRNAs encoding linker proteins of the phycobilisome, such as *apcE, cpcC1* and *cpcL*, as well as a protein of the ATP synthase complex that locates to the same membrane system. It is not unlikely that thylakoid membrane interactions of these proteins follow similar principles. This is in line with recent observations in an *Nostoc* sp. PCC 7120 deletion mutant lacking RbpG, the ortholog of *Synechocystis* 6803 Rbp3, which showed perturbed thylakoid membrane organization and function (58).

However, some mRNAs enriched by CLIP-Seq, such as *epsB*, encode proteins that are more likely to interact with the plasma membrane and even non-membrane proteins, most notably carboxysome shell proteins and RuBisCO (*rbcLXS* operon, see ***SI Appendix,* Fig. S5F**). These findings suggest a broader role for Rbp3, while supporting its previously proposed function, along with Rbp2, in localizing specific photosynthetic mRNAs to the thylakoid surface (30).

As interacting sites within the bound transcripts, we found mainly predicted folded secondary structure elements in which the RRMscorer algorithm identified U-rich pentanucleotide motifs (UUNGN) with at least one unpaired nucleotide or U-G base pair. Eukaryotic RRM proteins are highly flexible, typically binding single-stranded RNA, and occasionally single-stranded DNA (59, 60), although structured RNA elements are also documented (13). It is interesting to note that sequences within mRNA 3′UTRs have been recognized to contribute to the correct targeting of the encoded proteins to intracellular membranes (61, 62). The term memRBPs has been proposed for the “membrane-associated RNA-binding proteins” involved (63), and well-characterized memRBPs, like the fungal Rrm4, contain RRM domains (64). The analogy between Rbp3, which interacts with the 3′UTRs of mRNAs encoding key proteins of the photosynthetic machinery, and eukaryotic RRM proteins, which are involved in transport by targeting stem-loop elements in mRNA 3′UTRs, is striking. Thus, Rbp3 can be considered as a bacterial memRBP.

To perform their transport functions, memRBPs must interact with other cellular structures; for instance, Rrm4 shuttles along microtubules (64, 65). Although several elements of these mRNA transport systems have been elucidated, they are far from being fully understood (66).

Rbp3 likely does not function independently of other proteins. Mechanisms must exist to sort and recognize the targeted membrane or cellular compartment and to prevent premature translation. Therefore, we enriched proteins by co-IP to identify those interacting directly with Rbp3 or forming larger complexes nearby. A clear result of this approach was the enrichment of the entire set of ribosomal proteins (**Fig. 4**), indicating that complete ribosomes, not just subunits, were near Rbp3. However, recent fractionation of ribonucleoprotein complexes by Grad-Seq did not indicate co-fractionation of Rbp3 and ribosomes (7, 8). Thus, while a direct association of Rbp3 with ribosomes cannot be definitively excluded, there is no direct evidence supporting it, leading to the model shown in **Fig. 6A** in which Rbp3 is linked to ribosomes via a bound mRNA molecule.

**Fig. 6.**
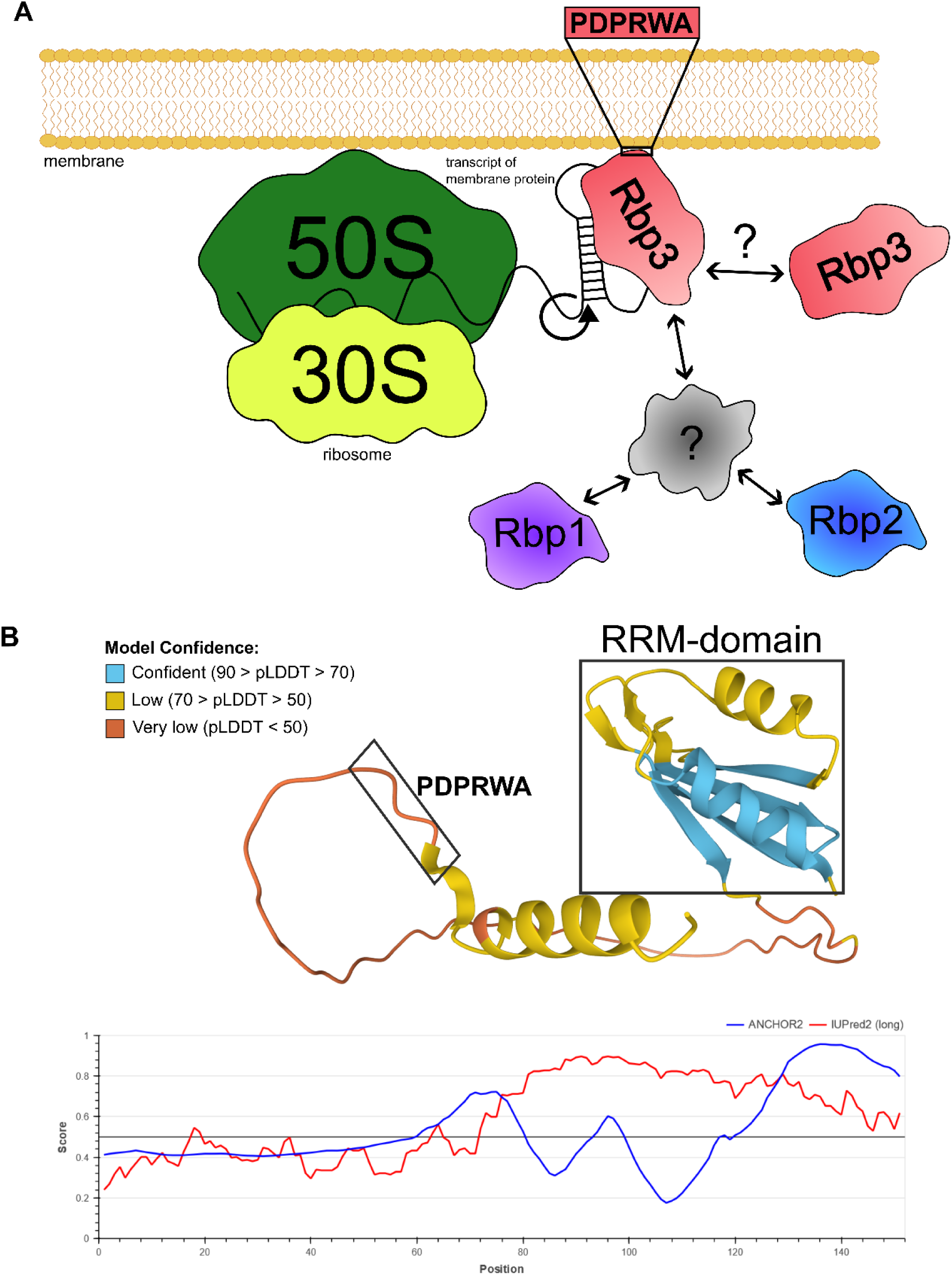
Proposed model of transcript localization towards the membrane mediated by Rbp3. **(A)** Rbp3 binds distinct mRNA motifs such as a secondary structure overlapping the stop codon in case of the *psaAB* mRNA. The Rbp3 interaction with the membrane surface might involve a PDPRWA sequence motif conserved within the C-terminal section of Rbp3 orthologs (26), identified by the DisoLipPred algorithm (67) as a lipid binding site. These mRNAs are translated near the membrane surface, explaining the high co-enrichment of ribosomal proteins with Rbp3. The helicase activity of the ribosome may unwind the RNA secondary structure, releasing Rbp3. Our data indicate that Rbp3 exists in both RNA-binding and non-binding states. The place where RNA binding is initiated and the factors involved in intracellular transport are unknown (question marks). The other RRM proteins, Rbp2 and Rbp1, likely also bind mRNAs with specific, potentially overlapping targets with Rbp3. Evidence suggests cross-talk among these Rbps, though its mechanisms are unclear. **(B)** Rbp3 structure predicted by AlphaFold (97, 98). The RRM domain characterized by a four-stranded antiparallel beta-sheet and two alpha helices, is boxed. The structure ends with a C-terminal short beta fold followed by an alpha helix. Between the RRM domain and the C-terminal structures, a long intrinsically disordered region is predicted, at the end of which is the PDPRWA element. The reliability of the prediction is color-coded; the intrinsically disordered region is confirmed by the AIUPred algorithm (99).

For mRNAs encoding PS core subunits, Rbp3 interacts with regions close to, or overlapping the stop codons. These mRNAs are highly translated and associated with polysomes, explaining the enrichment of ribosomal subunits in the Rbp3 co-IP. When the first ribosomes reach the stop codon region, their helicase activity likely unwinds the secondary structures where Rbp3 resides. As a result, it may be released and initiate interaction with another mRNA. Nascent polypeptides probably begin inserting into the membrane, keeping the mRNA-ribosome complexes associated with thylakoid surfaces, even after Rbp3 is released from the 3′ region. However, somehow membrane contact must be established and maintained. A conserved PDPRWA sequence motif within the C-terminal region of Rbp3 orthologs (26) is located within a large intrinsically disordered region of the protein (**Fig. 6B**). Prediction of putative lipid binding sites using the DisoLipPred algorithm, which was also trained on bacterial datasets (67), yielded a high score, suggesting its possible involvement in the interaction of Rbp3 with the membrane surface.

An unexpected finding in our co-IP experiments was the enrichment of Slr1870 and Slr0650, two structurally similar proteins with a LabA/NYN domain and an additional OST-HTH domain in Slr1870. A third such protein, Slr0755, was not enriched. The OST-HTH domain was predicted as RNA-binding and some eukaryotic proteins with that domain nucleate or organize ribonucleoprotein complexes (68). The relationship between Rbp3 and these proteins is unclear, but a similar structure is found in PIN-related RNases, which may be involved in tRNA or rRNA maturation. These proteins belong to the low-amplitude and bright (LabA) superfamily, including the *Synechococcus elongatus* LabA, which is linked to circadian timing (69). A LabA paralog, LalA, was described in *S. elongatus*, which is also involved in circadian gene expression, as well as in cell growth regulation (70). Recently, the *S. elongatus* Rbp2 homolog was reported to interact with the circadian clock protein KaiC, and disruption of its gene resulted in altered gene expression rhythms and KaiC localization (31). This suggests a connection between the cyanobacterial Rbps and the circadian clock system; however, Slr0755, the ortholog of *S. elongatus* LabA was not enriched in our co-IP, and the KaiC-interacting protein was the *S. elongatus* Rpb2 homolog and not Rbp3 addressed here.

Another interesting interacting protein was Slr0006, which binds tightly to membranes (71) and colocalizes with 30S ribosomal proteins and rRNA (72). Slr0006 has a TsaC fold, indicating its role in the N6-threonylcarbamoyladenosine modification in most ANN-decoding tRNAs (46), aligning with the enrichment of ribosomal proteins and other RNA maturation factors.

There is evidence of cross-talk between the three Rbps in *Synechocystis* 6803, as indicated by the upregulation of *rbp1* in the Δ*rbp2*Δ*rbp3* double mutant (**Fig. 5C, E, F**) and our observation that it is possible to generate double deletion mutants between any combination of these *rbp* genes, but not triple deletions. How this cross-talk is mediated is currently unknown, but we noticed the co-IP of possible regulatory proteins with Rbp3 such as Sll0518 (**Fig. 4A**). The gene *sll0518* is located directly downstream of *sll0517* encoding Rbp1, making it a candidate for possible cross-talk between different Rbps, especially during cold stress acclimation.

Similar targeting mechanisms probably exist for plasma membrane proteins (73) and, as our data indicate, potentially for cellular microcompartments, like carboxysomes. Moreover, the complex mechanisms involved in mRNA targeting may have contributed to the retention of genes in plant chloroplasts over evolutionary time scales (the ‘mRNA targeting hypothesis’) (74). While some components must be distinct, others might be shared. Clarification of the functions of the here identified non-ribosomal proteins and of the transcript segments that distinguish them from other transcripts is a promising topic for future research.

## Materials and Methods

### Culturing conditions and strain construction

*Synechocystis* 6803 GT-I (75) wild type and mutant strains were cultivated in copper-free BG11 supplemented with 20 mM TES buffer pH 7.5, under continuous white light of 50 µmol photons m^-2^ s^-1^ at 30°C (standard conditions). Mutant strains containing pVZ322 were cultivated in the presence of 50 µg/mL kanamycin and 2.5 µg/mL gentamycin, while Δ*rbp3* was kept at 50 µg/mL kanamycin. The Δ*rbp2*Δ*rbp3* double mutant strain was kept at 10 µg/mL chloramphenicol and 50 µg/mL kanamycin in the pre-cultures. Antibiotics were omitted during experimental assays. To test the acclimation responses of the Δ*rbp2*Δ*rbp3* double mutant and wild type to downshifts in temperature or to upshifts in light intensity, cultures were transferred at an OD_750_ of ∼ 1.0 from 30°C to 20°C, or at an OD_750_ of ∼0.6 from a light intensity of 50 to 250 µmol photons m^-2^ s^-1^. Kanamycin was omitted during co-IP experiments. For mRNA-FISH and Rbp3-GFP localization microscopy, *Synechocystis* 6803 PCC-M (76) was used due its larger cell size, facilitating high-resolution microscopy and protein localization. For details of the mutant strain construction and spectrometric characterization, see ***SI Appendix*.**

### Spectroscopic analyses

*Synechocystis* 6803 cultures for spectroscopic analyses were grown as described above but at light intensities of 20 µmol photons m^-2^ s^-1^. Room-temperature absorption spectra of cell suspensions were measured with a modernized Aminco DW-2000 ultraviolet-visible light spectrophotometer (Olis) and normalized to 750 nm. 77 K fluorescence emission spectra were recorded with a Perkin-Elmer LS55 luminescence spectrometer equipped with a liquid nitrogen housing. Cells were collected and prepared as previously reported (30). Fluorescence spectra were recorded with excitation at 435 nm or 600 nm and emission at 620–750 nm. Excitation and emission slit widths were 5 nm. Emission spectra were corrected for the instrument spectral response and normalized to the PSI or phycobilin fluorescence peaks after subtraction of the background signal.

### Protein induction, cell harvest and lysis

The expression of *rbp3-3xFLAG* as well as *sfGFP-3xFLAG* was induced in corresponding culture volumes at an OD_750_ of ∼ 0.6 by adding Cu_2_SO_4_ to a final concentration of 0.31 μM. After another 24 h of cultivation, the cells were harvested by centrifugation (4,000 *x* g, 4°C, 20 min). Cell pellet is resuspended in the respective lysis buffer. For cell lysis, mechanical disruption using a prechilled Precellys homogenizer (Bertin Technologies) was used. To remove intact cells and glass beads, the culture was centrifuged (800 *x* g, 5 min, 4°C), and the supernatant was collected for further processing. To solubilize membrane proteins, the cell lysates were incubated for 1 h in the presence of 2% n-dodecyl β-D-maltoside in the dark at 4°C. Afterwards, cell debris was removed by centrifugation (21,000 *x* g, 4°C 1 h). Supernatant was then used respectively.

### Production of recombinant Rbp3

To obtain the intact recombinant Rbp3 protein, it was first expressed as a GST fusion protein (GST-Rbp3) and then the GST tag was removed. The *rbp3* gene (*slr0193*) was amplified and ligated into vector pGEX-6P-1 using the In-Fusion Cloning kit (TaKaRa, Shiga, Japan) and the PCR primers pGEX-6P-1_inv-f/-r and Rbp3_iVEC-f/-r (***SI Appendix,* Table S5**). After verification of the sequences, the plasmids were introduced into *E. coli* Rosetta competent cells (TaKaRa) and used for the expression and purification of recombinant GST-Rbp3 protein as follows. The resulting *E. coli* strain was grown in 2 L of LB medium at 37°C. When the culture reached an OD_600_ of 0.6, IPTG was added to the medium at a final concentration of 1 mM and the growth temperature was shifted to 15°C. After 24 h, cells were harvested by centrifugation, washed with purification buffer (10% w/v glycerol,100 mM NaCl,20 mM HEPES-NaOH), and stored at −80°C until use. For protein purification, frozen cells were suspended in 200 mL of purification buffer. The cells were disrupted by sonication and centrifugated at 10,460 *× g* for 30 min at 4°C, and protein affinity purification was performed using an ÄKTA start chromatography system with a GSTrap column (Cytiva, MA, USA) according to the manufacturer’s instructions. The protein was eluted with 50 mM reduced glutathione and dialyzed against the dialysis buffer 1 (100 mM HEPES-KOH (pH 7.6), 100 mM NaCl, 1 mM 1,4-dithiothreitol (DTT), dialysis buffer 2 (50% w/v dialysis buffer 1, 50% w/v glycerol). Following dialysis, GST-Rbp3 was cleaved by PreScission Protease (Cytiva) to remove the GST-tag and dialyzed again. The purity of the protein was confirmed by SDS–PAGE and used for the BLI analysis.

### SDS-PAGE and Western blotting

Protein separation was done on by SDS-PAGE and protein detection with a subsequent western blot as described before (77). In short, after co-IP beads in elution buffer were mixed in a 5:1 ration with 5x protein loading dye (25 mM Tris-HCl pH 6.8, 25% glycerol, 10% SDS, 50 mM DTT, 0.05% bromophenol blue) and boiled for 5 min at 95°C. Equal volumes of supernatant were separated on a 15% SDS-PAA gel (78). After separation the proteins were transferred to a nitrocellulose membrane (Hybond™-ECL, Cytiva) at 1.5 A cm^-2^ for 1 h. After transfer, the membrane was blocked in 5% skimmed milk powder in TBS-T (50 mM Tris-HCl, 150 mM NaCl, 0.05% Tween20, pH 7.5) overnight at 4°C or for 1 h at room temperature. After blocking, the membrane was washed three times with TBS-T before incubating with mouse monoclonal anti-FLAG coupled to HRP (#A8592, Sigma) at a 1:5,000 titer in TBS-T. After incubation for 1 h at room temperature, signals were visualize using Western-Blot ECL spray (Advanta) and the FUSION SL Transilluminator (Vilber Lourmat).

### RNA preparation, Northern blotting and hybridization

Culture for RNA extraction was harvested by vacuum filtration on a hydrophilic polyether sulfone filter (Pall Supor-800) and snap frozen in liquid nitrogen. For RNA extraction, 1 mL PGTX (79) was given to the filter followed by an incubation of 15 min at 65°C. RNA was washed and precipitated as described previously (80).

The RNA concentration was determined by a Nanodrop spectrophotometer (peqlab). Equal amounts of RNA were diluted in water and 1:1 mixed with 2x RNA Gel Loading Dye (Thermo). Afterwards the samples were denatured at 72°C for 5 min.

RNA was separated on a 1.5% formaldehyde-agarose gel and RNA transfer to a positively charged nylon membrane (Amersham Hybond-N+, GE Healthcare) occurred overnight using a capillary blot system. Synthesis of probes and detection of radioactive signal has been described before (77).

### Microarray analysis

The here used Agilent microarrays have been designed for the direct detection of transcript after chemical labeling, without cDNA synthesis (81). The microarrays contain oligonucleotide probes for all annotated mRNAs and the majority of non-coding RNAs. Total RNA (5 µg) extracted from the wild type and deletion mutants grown at standard temperature (30°C) and after 1 h of cold stress (20°C) or after 0.5 h at high light was directly labeled using the ULS labeling kit for Agilent gene expression arrays with Cy3 according to the manufacturer’s protocol for Agilent one color microarrays (Kreatech Diagnostics, B.V., Netherlands). Array analysis was performed using two biological replicates and an internal technical replicate. Raw data were normexp background corrected (82) and quantile normalized using the limma R package (83). The transcriptome differences were calculated between wild type and mutant for the two different temperatures and the two different light conditions. Significance criteria for differential expression were log_2_FC ≥│1│, adj. p-value ≤ 0.05. P-values were adjusted for multiple testing with the Benjamini-Hochberg method. The array data has been deposited in the GEO database under the accession number GSE269515.

### Protein preparation and proteomic sample preparation

After harvesting, the cell pellet was resuspended in 1.5 mL FLAG-MS buffer (50 mM HEPES/NaOH pH 7, 5 mM MgCl_2_, 25 mM CaCl_2_, 150 mM NaCl) containing protease inhibitor (c0mplete, Roche). The cells were lysed as described above. Co-IP was performed by incubating 50 µL packed volume of Anti-FLAG M2 magnetic beads (Sigma) with the cleared cell lysate (1 h, 4°C, rotating). Afterwards the supernatant was removed using the DynaMag™-2 Magnet rack (Thermo Fisher Scientific) and the magnetic beads were washed 6x with FLAG-MS buffer. For additional details of the proteomic sample preparation and analyses by MS, see ***SI Appendix***.

### Crosslinking immunoprecipitation and sequencing (CLIP-Seq)

CLIP-Seq was carried out as described (10). Rbp3-3×FLAG and sfGFP-3×FLAG expression was induced in 250 mL *Synechocystis* cultures with 0.31 µM Cu_2_SO_4_ at an OD_750_ of 0.6-0.7. After 24 h, the cultures were transferred to 21 x14.5 x 5.5 cm^3^ plastic trays and crosslinked in a UV Stratalinker 2400 (Stratagene) three times at 0.45 J/cm^-2^. 25 mL culture aliquots were collected to prepare total RNA using the hot-phenol extraction method (80). The remaining culture was collected as described above. Cell pellets were resuspended in 1 mL lysis buffer (20 mM Tris/HCl pH 8.0, 1 mM MgCl_2_, 150 mM KCl, 1 mM DTT) with 1 U of RNase inhibitor per sample (RiboLock, Thermo Fisher Scientific) and lysed as described above. Supernatants were co-immunoprecipitated by incubation with 50 µL packed bead volume of Anti-FLAG M2 magnetic beads (Sigma) for 1 h at 4°C.The DynaMag™-2 magnetic rack (Thermo Fisher Scientific) was used for flow-through removal and washing. Beads were washed twice with FLAG buffer, then twice with high salt FLAG buffer (50 mM HEPES/NaOH pH 7, 5 mM MgCl_2_, 25 mM CaCl_2_, 1 M NaCl, 10% glycerol, 0.1% Tween20), and then washed two more times with FLAG buffer. Beads were subjected to proteinase K digestion (1% SDS, 10 mM Tris-HCl pH 7.5, 100 µg/mL proteinase K) for 30 min and 650 rpm at 50°C. Then, 200 µL PGTX (79) was added and the samples were boiled for 15 min at 65°C to extract protein-bound RNA. After adding 140 µL chloroform/isoamyl alcohol (24:1), the samples were incubated for 10 min at room temperature with agitation. For phase separation, the samples were centrifugated in a swing-out rotor at 3,260 x g for 3 min at room temperature. The upper aqueous phase was transferred to an RNase-free tube and mixed in a 1:1 ratio with 100% EtOH and processed via the clean and concentrator kit (Zymo Research). RNA was eluted in 15 µL RNase-free ddH_2_O. Construction of cDNA libraries and sequence analysis were performed by a commercial provider (Vertis Biotechnologie AG Freising, Germany). The RNA samples were fragmented using ultrasound (4 pulses of 30 s each at 4°C). An oligonucleotide adapter was ligated to the RNA 3’ ends. M-MLV reverse transcriptase and a 3′ primer complementary to the adapter sequence were used for first-strand cDNA synthesis. The obtained cDNA was purified and the 5’ Illumina TruSeq sequencing adapter ligated to the cDNA 3’ ends. The cDNA was amplified by 13 cycles of PCR (15 to 18 for the negative control), to about 10-20 ng/μL using a high-fidelity DNA polymerase and introducing TruSeq barcode sequences as part of the 5’ and 3’ TruSeq sequencing adapters. The cDNA was purified using the Agencourt AMPure XP kit (Beckman Coulter Genomics) and analyzed by capillary electrophoresis. For Illumina NextSeq sequencing, the samples were pooled in approximately equimolar amounts. The cDNA pool was 200 – 600 bp size-fractionated via a preparative agarose gel and sequenced on an Illumina NextSeq 500 system with a read length of 75 bp.

Raw reads of the RNA-seq libraries were analyzed using the galaxy platform. The galaxy workflow can be accessed and reproduced at the following link: https://usegalaxy.eu/u/luisa_hemm/w/poi-3xflag-clip. In short, read quality was check with FastQC in the beginning. After the quality check, reads were mapped to the *Synechocystis* chromosome (GenBank accession NC_000911.1) and the 4 large plasmids pSYSA, pSYSG, pSYSM and pSYSX (NC_005230.1, NC_005231.1, NC_005229.1, NC_005232.1) using BWA-MEM (84) with default parameters. Resulting data were filtered using the BAM filter (85) to remove unmapped reads as well as reads smaller than 20 nucleotides. The annotation gff-file was based on the reference GenBank annotations complemented by additional genes based on transcriptomic analyses (86). htseq-count (87) (mode union, nonunique all, minaqual 0) was used to count the reads per transcript. Only transcripts with ≥ 10 reads in at least three samples were subjected to differential expression analysis with DESeq2 (36). To visualize coverage of the reads, bam files were transformed into wiggle files using RSeQC (88) and summarizing the reads from the triplicate analyses for each position.

### PNK assay

Procedures followed previously described methods (8, 35). Rbp3-3*×*FLAG and sfGFP- 3*×*FLAG expression was induced in 250 mL *Synechocystis* cultures with 0.31 µM Cu_2_SO_4_ at an OD_750_ of 0.6–0.7. After 24 h, cultures were transferred to 21 x14.5 x 5.5 cm^3^ plastic trays on ice and crosslinked three times at 0.75 J/cm^-2^ using a UV Stratalinker 2400 (Stratagene) to induce protein-RNA covalent bonds. Negative controls were not crosslinked. Cells were collected by centrifugation and the pellets were resuspended in 800 µL of NT-P buffer (50 mM NaH_2_PO_4_, 300 mM NaCl, 0.05% Tween, pH 8.0) containing c0mplete protease inhibitor (Roche). Cell lysis supernatants were co-immunoprecipitated by incubation with 80 µL packed bead volume of Anti-FLAG M2 magnetic beads (Sigma) for 1 h at 4°C with slight rotation. The DynaMag™- 2 magnetic rack (Thermo Fisher Scientific) was used for flow-through removal and washing using 2 mL per wash step. Beads were washed twice with high salt NT-P buffer (50 mM NaH_2_PO_4_, 1 M NaCl, 0.05% Tween, pH 8.0), once with NT-P buffer and once with benzonase buffer (50 mM Tris-HCl, 1 mM MgCl_2_, pH 8). Beads were resuspended in 100 µL benzonase buffer containing 25 U benzonase (Sigma). After incubation for 10 min at 37°C and 900 rpm, beads were washed once with high-salt NT-P buffer and twice with FastAP buffer (10 mM Tris-HCl, 5 mM MgCl_2_, 100 mM KCl, pH 8). Beads were then resuspended in 150 µL FastAP buffer and incubated with 2 U Fast-AP (Thermo) for 30 min at 37°C and 900 rpm. Beads were washed once with high-salt NT-P buffer and twice with PNK buffer (50 mM Tris-HCl, 10 mM MgCl2, 0.1 mM spermidine, pH 8) and then resuspended in 100 µL PNK buffer containing 10 µCi of [γ-^32^P]-ATP and 1 µL PNK (Thermo). After incubation at 37°C for 30 min at 900 rpm beads were washed once with high salt NT-P buffer and twice with NT-P buffer. For elution, beads were resuspended in 35 µL TBS (50 mM Tris-HCl, 150 mM NaCl, pH 7.5) with 0.5 µg/µL FLAG peptide (Sigma) and incubated at room temperature for 15 min at 900 rpm. To the supernatant 5 µL protein loading dye (25 mM Tris-HCl pH 6.8, 25% glycerol, 10% SDS, 50 mM DTT, 0.05% bromophenol blue) was added and samples were boiled for 5 min at 95°C. Proteins were separated by SDS-PAGE and western blotting was done as described above. After transfer to the membrane, the membrane was exposed for several hours to a phosphoimaging screen (Kodak) and visualized using a Typhoon FLA 9500 (GE Healthcare), followed by western blot incubation.

### Bio-Layer Interferometry

Recombinant Rbp3 protein interactions with RNA oligonucleotide ligands were examined using the BLI system, BLItz (Fortebio). Ligand RNA oligonucleotides were synthesized with a biotinylated 5’ end (Eurofins) and used for BLI analysis. The binding buffer (4% glycerol, 10 mM Tris-HCl pH 7.5, 1 mM MgCl₂, 0.5 mM EDTA, 50 mM NaCl, 0.5 mM DTT, 0.05 mg/mL poly(dI-dC)·poly(dI-dC), 0.05% [w/v] BSA, 0.01% Tween 20) was used to dilute both ligand RNA and analyte Rbp3 protein. Experiments were conducted at room temperature. 0.5 pmol of the biotinylated ligand RNA oligonucleotides were captured on a streptavidin biosensor (Sartorius) that had been equilibrated with binding buffer. Rbp3 was bound to the ligand RNA-reacted biosensor proteins at concentrations of 0.5, 1, 2 or 3 pmol/μL, and the association was analyzed. The same binding buffer was used for the dissociation step.

### mRNA fluorescent *in situ* hybridization

mRNA-FISH was performed as described (89), with adjustments for *Synechocystis* 6803 according to reference (30). Briefly, the target RNA was hybridized with 44–48 oligonucleotides, each 20 nt long, fused to a TAMRA fluorophore at the 3’ end. Oligonucleotide probes for mRNA-FISH were sourced from LGC Biosearch Technologies (***SI Appendix,* Table S6)**. For further procedural details, including cell fixation, permeabilization, hybridization and buffer compositions, see reference (30). For imaging hybridized *Synechocystis* 6803 PCC-M cells, fixed samples were mounted on a high precision cover glass covered by a 1.5% agarose pad to minimize cell movement. Imaging was conducted using an LSM 880 confocal microscope equipped with an Airyscan detector (Carl Zeiss) and a Plan-Apochromat 63×/1.4NA oil objective lens. Images were recorded in 16-bit format, at 488 × 488 pixels resolution, 8x line averaging and a zoom value of 10. The 561 nm, 488 nm, and 633 nm lasers were used to excite the TAMRA fluorophore, Rbp3-sfGFP fusion protein, and thylakoid membrane photosynthetic pigments, respectively. To prevent signal bleed-through, a secondary beam splitter and filter set were installed before the Airyscan detector. A short pass 615 nm beam splitter specified TAMRA emission, and a band pass filter from 495–550 nm isolated sfGFP emission. Image reconstruction used the Airyscan processing software with automatic strength.

Prior to in-depth image analysis, autofluorescence from RNA-FISH and Rbp3-sfGFP was removed by background subtraction as described (30). The ImageJ plugin EzColocalization was used to determine the cell area, calculate mean fluorescence intensity and perform colocalization analysis (90, 91). Binary images of cell outlines were created from the thylakoid membrane channel using the auto threshold command. The adjustable watershed with a tolerance of 5 was applied to distinguish between adjacent cells. Cells not fully captured in the images were excluded from analysis. Colocalization of hybridized target mRNA and Rbp3-sfGFP fluorescence was assessed by calculating the Pearson correlation coefficient using Costes thresholding (92).

## Supporting information

Supplemental Figures, Tables and Overview

## Data availability

The CLIP-seq data have been deposited in the SRA database under the accession number PRJNA1104562. The mass spectrometry proteomics data have been deposited to the ProteomeXchange Consortium (93) via the PRIDE (94) partner repository with the dataset identifier PXD045584. All mass spectrometry proteomics datasets analyzed during this study are available online at the MassIVE repository (http://massive.ucsd.edu/; dataset identifier: MSV000092937). The microarray data have been deposited in the GEO database under the accession number GSE269515.

## Author Contributions

WRH designed the project and secured funding. ST and OS carried out MS-based proteomic analyses. SW, SM and KK provided the Δ*rbp2/3* mutant and the BLI data. VR performed the microarray analyses, MM and CWM provided the spectroscopic analyses. EL and AW performed the RNA-FISH and microscopical analyses and contributed to the data interpretation. All other experiments and analyses were performed by LH. LH and WRH wrote the manuscript with input from all authors.

## Classification

Biological Sciences, Microbiology

