## Supplemental Figures, Tables and Overview for "The RRM domain-containing protein Rbp3 interacts with ribosomes and the 3′ ends of mRNAs encoding photosynthesis proteins"

### Supplementary Information

**Supplementary Materials and Methods:** p. 2
**Overview on Supplementary Datasets** p. 6
**Supplementary Tables:** p. 7
**Supplementary Figures:** p. 22
**Supplementary References:** p. 28

### Supplementary Materials and Methods

#### Construction of mutant and tagged strains

*Synechocystis* 6803 GT-I (1) served as the wild type and background for constructing Δ*rbp3* and Δ*rbp2*Δ*rbp3* mutants (2). Strains harboring the pVZ322 plasmid with the *rbp3-3xFLAG* gene under control of the *petE* promoter ($P_{petE}$) were previously described (2). To construct the sfGFP-3xFLAG expression plasmid, the *sfGFP* sequence with overlaps to $P_{petE}$ was PCR-amplified from the pXG10-SF plasmid (3) (primer and plasmid details in ***SI Appendix,*** **Table S5**). For subcloning, pJet1.2-P*petE*-*slr0193* (2) was used, replacing the *slr0193* sequence with *sfGFP* using AQUA cloning. The constructed vector was subjected to inverse PCR to introduce the 3xFLAG tag to the C-terminus of sfGFP followed by *Dpn*I digest (Thermo Fisher Scientific), 5′ phosphorylation (T4 PNK, Thermo Fisher Scientific), self-ligation (T4 DNA ligase, Thermo Fisher Scientific) and heat-shock transformation into *E. coli* DH5α. The $P_{petE}$-*sfGFP*-3xFLAG fragment was amplified and inserted by AQUA cloning into the self-replicating, multi-host vector pVZ322 linearized by restriction with *Hind*III and *Xho*I. To obtain the complementation strain with the pVZ322::*rbp3* plasmid, the *rbp3* gene was amplified from genomic DNA including the native promoter and 3′UTR using specific primers (***SI Appendix,*** **Table S5**). Cloning into pVZ322 and transformation of *Synechocystis* 6803 GT-I was done as described for the strain expressing the triple FLAG-tagged Rbp3 (2).

To generate *rbp3* knockouts in the PCC-M strain, total gDNA of Δ*rbp3* GT-I cells containing the *rbp3* knockout cassette was used to transform PCC-M wild-type cells. Transformants were selected on kanamycin (50 µg/mL) containing BG11 plates. Complete segregation was confirmed by colony PCR (primers FP and RP, ***SI Appendix,*** **Table S5**).

To generate PCC-M Δ*rbp3* + $P_{rbp3}$-Rbp3-sfGFP strains, the *rbp3* knockout strain was transformed with the pVZ322 plasmid containing the *rbp3-sfGFP* gene under the control of the native *rbp3* promoter ($P_{rbp3}$). To obtain the sfGFP-tagged version of Rbp3, the 3xFLAG fragment in the *rbp3-3xFLAG* construct was replaced by the *sfGFP* gene by AQUA cloning. Successful exchange of the 3xFLAG segment by the *sfGFP* gene was confirmed by PCR and by Sanger sequencing (GATC, Eurofins). Deletion of *rbp3* in PCC-M background led to smaller cells and a reduced chlorophyll content similar to the *rbp3* mutant in the GT-I strain, a phenotype which could be complemented by

Rbp3-GFP indicating that the GFP fusion did not interfere with the functionality (***SI Appendix,*** **Fig. S3**).

**Spectrometric measurements**

Whole-cell absorption spectra were obtained using an SLM Aminco DW-2000 spectrophotometer. Cyanobacteria were grown to mid-log phase in BG11 medium supplemented with 20 mM TES buffer pH 7.5 under continuous white light at 15-20 µmol photons $m^{-2}$ $s^{-1}$ at 30°C. Spectra were recorded from 400 nm to 750 nm and normalised at 750 nm. Data presented are the representative of similar results from 3 independent cultures. Fluorescence emission spectra were recorded at 77 K with a Perkin Elmer LS55 fluorescence spectrometer with a liquid nitrogen housing. Cultures were grown as for the absorption spectra, concentrated to 5 µM chlorophyll in BG11 medium and loaded into silica capillary tubes. The tubes were dark-adapted for 10 min and then snap frozen in liquid nitrogen. Fluorescence emission spectra were recorded from 620-750 nm with slit-width 3-5 nm and excitation at 435 nm for chlorophyll *a* or at 600 nm for phycocyanin. Spectra were normalized to the PSI peak (~735 nm) or to the phycocyanin/allophycocyanin peak (~660 nm) and are representative of 3 independent and similar results.

**Proteomic sample preparation and analyses by MS**

Proteins were further processed according to the manufacturer's protocol for S-Trap micro filters (Protifi), and using the provided buffers. Briefly, 60 µL of the 1x lysis buffer (5% SDS, 50 mM TEAB pH 8.5) was added to the beads and incubated at 95°C for 10 min to elute the proteins. In the following process 46 µL of eluate were used. Proteins were reduced by incubation with reduction buffer (end concentration 5 mM Tris(2-carboxyethyl)phosphine (TCEP)) for 15 min at 55°C and alkylated by incubation with alkylation buffer (end concentration 20 mM methyl methanethiosulfonate (MMTS)) for 10 min at room temperature in the dark. Afterwards, a final concentration of 1.2% phosphoric acid and then six volumes of binding buffer (90% methanol; 100 mM TEAB; pH 7.55) were added. After gentle mixing, the protein solution was loaded to an S-Trap filter and spun 3 times at 10,000 *x* g for 30 sec. The filter was washed three times using 150 µL of binding buffer. Sequencing-grade trypsin (Promega, 1:25 enzyme:protein ratio) diluted in 20 µL digestion buffer (50 mM TEAB) were added to the filter and digested at 47°C for 2 h. To elute peptides, three step-wise buffers were applied:

a) 40 µL 50 mM TEAB, b) 40 µl 0.2% formic acid in $H_2O$, and c) 50% acetonitrile and 0.2% formic acid in $H_2O$. Each step was eluted by centrifugation (10,000 *x* g for 1 min). The peptide solutions were combined and dried in a DNA SpeedVac DNA120 (Savant). For LC-MS/MS measurements 800 ng of peptides were analyzed on an LTQ-Orbitrap Elite mass spectrometer (Thermo Fisher Scientific) coupled to an EASY-nLCTM 1000 UHPLC system (Thermo Fisher Scientific). The column setup consisted of a Acclaim™ PepMap™ 100 C18 column (Thermo Fisher Scientific, Cat. No. 164946) and a 200 cm µPac GEN1 analytical column (PharmaFluidics, 55250315018210) coupled to a Nanospray FlexTM ion source (Thermo Scientific, ES071) and a fused silica emitter (MS Wil, TIP1002005-5).

For peptide separation, a linear gradient of buffer B (0.1% formic acid in 80% acetonitrile, Fluka) was applied, increasing from 8 to 55% over 80 min and from 55 to 100% over the next 40 min. Peptides were analyzed using data-dependent acquisition (DDA). In the positive ion mode, a full scan ranging from m/z 400 to 1600 with a resolution of 60,000 and a maximum ion accumulation time of 40 ms was performed. The top 15 precursors of the highest abundance in the full scan were selected and fragmented by collision-induced dissociation (CID) function and analyzed in MS/MS with a normalized collision energy of 35%, 10 ms activation time, and a minimum TIC threshold of 10,000 for MS/MS selection. The following dynamic exclusion settings included: repeat counts 1; repeat duration 15 s; exclusion duration 35 s.

Raw data were analyzed with MaxQuant (v 1.6.17.0) with the built-in Andromeda peptide search engine (4). The false discovery rate (FDR) at both the protein and peptide level was set to 1%. Two missed cleavage sites were allowed, no variable modifications were allowed, and carbamidomethylation of cysteines was set as fixed modification. The match between runs option was selected. For label free quantification the MaxLFQ algorithm was applied using the standard settings. Only unique peptides were used for quantification. The database "Synechocystis sp. (strain PCC 6803 / Kazusa)" was downloaded from https://www.uniprot.org/proteomes/UP000001425 on Jun 9th, 2022.

Data were normalized on peptide level by equalizing the medians using the MSstats package (v. 3.20.3) in R (v. 4.0.4) (5). Subsequently, protein LFQ intensities were $\log_2$ transformed. P-values were adjusted using the Benjamini-Hochberg procedure.

The proteome raw data acquired by MS were deposited at the ProteomeXchange Consortium (http://proteomecentral.proteomexchange.org) via the PRIDE partner repository (6) under the identifier PXD045584. The intensities were compared using LFQ (label-free quantification) values (7) with Perseus (version 1.6.1.3 (8)). In summary, contaminants, reverse sequences and proteins only identified by site were removed from the matrix and LFQ intensities were $log_2$-transformed. Before t-test and visualization using a volcano plot, the missing values were replaced by imputation with the normal distribution for each column separately (default settings). For hierarchical clustering (default parameters), only proteins with three valid values in at least one declared group (Rbp3_3×FLAG and sfGFP_3xFLAG) were considered.

**Overview on Supplementary Datasets**

**Dataset 1 "Cold".** Transcriptomic differences (microarray data) before and 1 h after transfer of *Synechocystis* 6803 wild type and Δ*rbp2*Δ*rbp3* double mutant cultures at an $OD_{750}$ of ~ 1.0 from 30°C to 20°C. Previously defined transcription units (TU) and the coverage from previous RNA-seq analyses of exponentially growing cultures (light grey) or after transfer to 15 °C (dark grey) are indicated by the coverage plots in the background, with the read counts plotted on the right X-axis (9).

**Dataset 2 "HL".** Transcriptomic differences (microarray data) before and 1 h after transfer of *Synechocystis* 6803 wild type and Δ*rbp2*Δ*rbp3* double mutant cultures at an $OD_{750}$ of ~ 0.6 from a light intensity of 50 to 250 µmol photons $m^{-2}$ $s^{-1}$. Previously defined transcription units (TU) and the coverage from previous RNA-seq analyses of exponentially growing cultures (light grey) or after transfer to high light (dark grey) are indicated by the coverage plots in the background, with the read counts plotted on the right X-axis ((9).

For both Datasets: The chromosome and all 7 plasmids are arranged in a linear fashion, microarray probes are indicated by short vertical tabs and the $log_2$ intensity values after hybridization are plotted on the left X-axis.
The raw data have also been deposited in the GEO database under the accession number GSE269515.

### Supplementary Tables

**Table S1. Transcripts significantly enriched in the co-IP with Rbp3 after UV cross-linking.** The locus tag is given, together with the enrichment ($log_2FC$), *p* value and adjusted *p* value (*p* adj) compared to the sfGFP negative control. The next columns give the gene name if known or a comment, followed by the classification of the respective protein as associated to thylakoid membrane (TM), plasma membrane (PM) or cell envelope (ENV). Cytoplasmic proteins (CP) and carboxysomal (CBX) proteins are also indicated. The localization is based on the existing annotation in the uniprot database (10) and considering published proteome mapping data (11). Unclear localization is indicated by a dash. Locus tags starting with ncl/ncr indicate previously defined non-coding transcripts (9). The underlying data are average read counts from three biological replicates each for the co-IP with Rbp3 and sfGFP, respectively. PBS, phycobilisome. This table extends the data in **Fig. 2C**.

| Locus tag | $log_2FC$ | *p* value | p adj | Gene | Localization |
|---|---|---|---|---|---|
| slr1835 | 3.87 | 4.01E-48 | 1.03E-44 | psaB | TM |
| slr1834 | 3.41 | 4.74E-43 | 8.10E-40 | psaA | TM |
| ncr1220 | 2.56 | 4.34E-19 | 1.39E-16 | 3’ petD | - |
| ssl2615 | 2.56 | 1.08E-27 | 1.11E-24 | atpH | TM |
| isiE | 2.54 | 7.01E-14 | 8.17E-12 | isiE | TM |
| slr1311 | 2.47 | 2.08E-20 | 9.68E-18 | psbA2 | TM |
| sll1867 | 2.41 | 6.75E-20 | 2.47E-17 | psbA3 | TM |
| slr0335 | 2.28 | 2.76E-27 | 2.36E-24 | apcE | TM |
| sll0923 | 2.26 | 1.15E-31 | 1.47E-28 | epsB | PM |
| slr0146 | 2.10 | 4.97E-16 | 9.10E-14 | PBS protein | TM |
| sll1471 | 2.06 | 3.73E-15 | 5.97E-13 | cpcL | TM |
| ssr2553 | 2.02 | 9.40E-19 | 2.84E-16 | ssr2553 | - |
| ncr0410 | 1.98 | 1.70E-05 | 0.000296 | 3’ slr1544 | - |
| sll1733 | 1.94 | 3.23E-22 | 1.84E-19 | ndhD3 | TM |
| sll0450 | 1.92 | 5.05E-23 | 3.24E-20 | norB | - |
| slr0373 | 1.91 | 6.45E-21 | 3.30E-18 | slr0373 | CP |
| sll1934 | 1.88 | 1.02E-11 | 8.07E-10 | sll1934 | - |
| sll1028 | 1.88 | 4.86E-18 | 1.31E-15 | ccmK2 | CBX |
| sll1580 | 1.87 | 4.20E-20 | 1.80E-17 | cpcC1 | TM |
| sll1070 | 1.87 | 1.27E-17 | 2.97E-15 | tktA | CP |
| ncr1020 | 1.86 | 2.18E-10 | 1.29E-08 | 5’ glnN | - |
| sll1324 | 1.84 | 1.69E-11 | 1.29E-09 | atpF | TM |
| slr0186 | 1.83 | 1.15E-16 | 2.18E-14 | leuA | CP |
| sll1639 | 1.82 | 4.41E-18 | 1.26E-15 | ureD | CP |
| slr1841 | 1.81 | 2.40E-17 | 5.13E-15 | porin | ENV |

| | | | | | |
|---|---|---|---|---|---|
| slr1986 | 1.80 | 4.77E-14 | 6.11E-12 | apcB | TM |
| sll1734 | 1.80 | 2.57E-13 | 2.75E-11 | cupA | TM |
| sll1732 | 1.79 | 3.79E-17 | 7.77E-15 | ndhF3 | TM |
| ncl0610 | 1.78 | 7.49E-07 | 1.93E-05 | 3’ sll1150 | - |
| sll1323 | 1.77 | 2.31E-14 | 3.49E-12 | atpG | TM |
| sll1579 | 1.76 | 5.22E-14 | 6.52E-12 | cpcC2 | TM |
| slr0954 | 1.75 | 2.67E-12 | 2.36E-10 | rnc_mini | CP |
| sll0760 | 1.75 | 8.35E-12 | 6.80E-10 | ycf38 | - |
| slr0601 | 1.73 | 1.26E-13 | 1.43E-11 | slr0601 | PM |
| slr0287 | 1.71 | 2.45E-13 | 2.67E-11 | khpA | - |
| ssr1552 | 1.70 | 3.81E-12 | 3.20E-10 | ssr1552 | - |
| slr0012 | 1.69 | 1.89E-11 | 1.43E-09 | rbcS | CBX |
| slr2051 | 1.68 | 4.05E-14 | 5.33E-12 | cpcG1 | TM |
| slr0374 | 1.68 | 2.26E-08 | 8.35E-07 | ycf46 | - |
| slr1516 | 1.66 | 9.63E-12 | 7.71E-10 | sodB | CP |
| slr1908 | 1.65 | 1.43E-13 | 1.59E-11 | porin | ENV |
| slr1856 | 1.64 | 1.49E-08 | 5.70E-07 | icfG | - |
| slr0929 | 1.63 | 1.87E-10 | 1.15E-08 | parA family | - |
| sll0928 | 1.63 | 2.91E-14 | 4.03E-12 | apcD | TM |
| sll0218 | 1.63 | 4.66E-08 | 1.67E-06 | sll0218 | - |
| slr0882 | 1.62 | 3.25E-10 | 1.81E-08 | ycf84 | PM |
| sll0108 | 1.62 | 1.60E-17 | 3.57E-15 | amt1 | PM |
| sll0851 | 1.62 | 1.83E-10 | 1.14E-08 | psbC | TM |
| slr0007 | 1.60 | 1.71E-14 | 2.65E-12 | slr0007 | - |
| slr1756 | 1.60 | 4.85E-10 | 2.61E-08 | glnA | CP |
| slr0147 | 1.59 | 3.33E-13 | 3.48E-11 | slr0147 | TM |
| sll1951 | 1.59 | 2.56E-11 | 1.88E-09 | sll1951 | PM |
| sll1577 | 1.58 | 6.59E-11 | 4.39E-09 | cpcB | TM |
| sll1950 | 1.58 | 5.80E-14 | 6.92E-12 | sll1950 | - |
| sll1304 | 1.58 | 1.42E-07 | 4.54E-06 | sll1304 | - |
| slr0011 | 1.58 | 3.72E-09 | 1.66E-07 | rbcX | CBX |
| slr1273 | 1.57 | 6.91E-10 | 3.58E-08 | slr1273 | - |
| sll1535 | 1.57 | 5.71E-14 | 6.92E-12 | rfbP | ENV |
| slr1281 | 1.56 | 5.27E-11 | 3.60E-09 | ndhJ | TM |
| slr0331 | 1.55 | 5.88E-18 | 1.51E-15 | ndhD1 | TM |
| slr1546 | 1.52 | 3.84E-13 | 3.94E-11 | slr1546 | - |
| sll1735 | 1.52 | 1.14E-08 | 4.66E-07 | cupS | TM |
| slr1655 | 1.49 | 5.20E-13 | 5.13E-11 | psaL | TM |
| slr0565 | 1.49 | 2.00E-10 | 1.20E-08 | SyndsbAB | PM |
| sll1578 | 1.48 | 1.48E-08 | 5.70E-07 | cpcA | TM |
| slr1127 | 1.48 | 2.88E-14 | 4.03E-12 | slr1127 | - |
| sll0272 | 1.48 | 1.68E-08 | 6.39E-07 | ndhV | - |
| sll1322 | 1.48 | 2.27E-09 | 1.07E-07 | atpI | TM |
| sll1099 | 1.48 | 5.14E-10 | 2.72E-08 | tufA | CP |
| sll1274 | 1.47 | 5.96E-09 | 2.53E-07 | sll1274 | - |

| | | | | | |
|---|---|---|---|---|---|
| slr1855 | 1.46 | 9.93E-13 | 9.16E-11 | slr1855 | - |
| sll1898 | 1.46 | 4.12E-06 | 8.70E-05 | sll1898 | - |
| slr0144 | 1.46 | 3.03E-08 | 1.11E-06 | slr0144 | - |
| sll0682 | 1.45 | 8.80E-06 | 0.0001658 | pstA | PM |
| sll1931 | 1.45 | 3.75E-12 | 3.20E-10 | glyA | CP |
| MYO_RS17880 | 1.45 | 9.85E-16 | 1.63E-13 | slr8038 | - |
| sll0223 | 1.44 | 7.36E-17 | 1.45E-14 | ndhB | TM |
| slr0009 | 1.44 | 2.47E-09 | 1.14E-07 | rbcL | CBX |
| slr1177 | 1.44 | 2.13E-11 | 1.58E-09 | slr1177 | PM |
| sll1338 | 1.44 | 1.00E-12 | 9.16E-11 | sll1338 | CP |
| sll0219 | 1.44 | 1.92E-08 | 7.17E-07 | flv2 | - |
| sll1665 | 1.44 | 2.66E-12 | 2.36E-10 | sll1665 | PM |
| sll0469 | 1.43 | 2.56E-14 | 3.75E-12 | prsA | CP |
| sll0757 | 1.42 | 8.79E-16 | 1.50E-13 | purF | CP |
| slr0737 | 1.42 | 2.21E-08 | 8.21E-07 | psaD | TM |
| ssr1600 | 1.42 | 2.17E-10 | 1.29E-08 | ssr1600 | CP |
| slr1599 | 1.41 | 1.96E-09 | 9.31E-08 | Z_ISO | - |
| slr0394 | 1.40 | 4.38E-09 | 1.91E-07 | pgk | CP |
| sll0819 | 1.40 | 4.89E-10 | 2.61E-08 | psaF | TM |
| slr1545 | 1.40 | 5.05E-13 | 5.08E-11 | sigG | - |
| slr1744 | 1.39 | 5.77E-13 | 5.58E-11 | amiA | PM |
| sll0418 | 1.39 | 1.00E-10 | 6.50E-09 | vte1 | - |
| sll1130 | 1.39 | 3.51E-06 | 7.64E-05 | sll1130 | CP |
| sll1091 | 1.38 | 5.91E-10 | 3.09E-08 | chlP | - |
| slr0906 | 1.38 | 9.73E-10 | 4.85E-08 | psbB | TM |
| sll1263 | 1.37 | 5.70E-12 | 4.72E-10 | sll1263 | PM |
| sll0018 | 1.37 | 9.73E-08 | 3.26E-06 | fbaA | CP |
| slr1459 | 1.37 | 8.21E-10 | 4.17E-08 | apcF | TM |
| slr1886 | 1.36 | 4.40E-09 | 1.91E-07 | slr1886 | CP |
| sll1325 | 1.36 | 9.16E-09 | 3.79E-07 | atpD | TM |
| slr0059 | 1.36 | 5.03E-05 | 0.0007451 | slr0059 | - |
| sll1327 | 1.36 | 3.02E-09 | 1.38E-07 | atpC | TM |
| sll0217 | 1.35 | 1.81E-06 | 4.23E-05 | flv4 | - |
| sll1785 | 1.34 | 2.05E-06 | 4.75E-05 | cucA | PM |
| slr0042 | 1.33 | 1.88E-10 | 1.15E-08 | porin | PM |
| ssl0410 | 1.32 | 1.43E-08 | 5.58E-07 | ssl0410 | - |
| sll0461 | 1.31 | 1.04E-11 | 8.11E-10 | proA | CP |
| slr1571 | 1.31 | 6.32E-06 | 0.0001246 | slr1571 | PM |
| slr0343 | 1.31 | 5.82E-09 | 2.49E-07 | petD | TM |
| slr0040 | 1.31 | 1.02E-05 | 0.0001903 | cmpA | PM |
| slr0993 | 1.30 | 1.25E-07 | 4.07E-06 | nlpD | CP |
| slr1708 | 1.30 | 1.42E-08 | 5.58E-07 | slr1708 | ENV |
| sll1488 | 1.30 | 2.48E-10 | 1.44E-08 | M23 peptidase | - |
| sll0849 | 1.30 | 5.92E-08 | 2.04E-06 | psbD | TM |
| sll0519 | 1.30 | 3.75E-11 | 2.71E-09 | ndhA | TM |

| | | | | | |
|---|---|---|---|---|---|
| slr0257 | 1.29 | 1.75E-08 | 6.60E-07 | ctpB | CP |
| slr1329 | 1.29 | 9.01E-09 | 3.75E-07 | atpB | TM |
| slr1137 | 1.28 | 1.01E-09 | 4.97E-08 | ctaDI, coxA | TM |
| sll0273 | 1.28 | 1.19E-09 | 5.80E-08 | nhaS2 | PM |
| sll1108 | 1.28 | 1.87E-07 | 5.89E-06 | surE | CP |
| slr0447 | 1.28 | 3.90E-07 | 1.10E-05 | urtA | PM |
| slr0164 | 1.27 | 2.03E-07 | 6.27E-06 | clpP4 | CP |
| slr0602 | 1.27 | 1.82E-06 | 4.23E-05 | slr0602 | - |
| slr1503 | 1.27 | 2.48E-09 | 1.14E-07 | slr1503 | - |
| slr0148 | 1.26 | 3.19E-07 | 9.35E-06 | fdx5 | - |
| slr0679 | 1.26 | 8.68E-10 | 4.36E-08 | rsmB | CP |
| smr0014 | 1.26 | 0.000698 | 0.006816 | smr0014 | - |
| sll7047 | 1.26 | 2.49E-10 | 1.44E-08 | sll7047 | PM |
| slr0006 | 1.25 | 3.37E-09 | 1.51E-07 | tsaC | CP |
| sll1783 | 1.25 | 0.0001463 | 0.0018424 | sll1783 | CP |
| MYO_RS16880 | 1.25 | 1.20E-06 | 2.95E-05 | sll5069 | - |
| sll0543 | 1.24 | 0.0014233 | 0.011943 | sll0543 | - |
| sll1886 | 1.23 | 7.32E-07 | 1.89E-05 | sll1886 | - |
| slr0358 | 1.23 | 4.78E-08 | 1.70E-06 | slr0358 | PM |
| sll1713 | 1.23 | 4.65E-08 | 1.67E-06 | hisC | - |
| sll0041 | 1.22 | 7.08E-10 | 3.63E-08 | pixJ1, taxD1 | PM |
| ncl0070 | 1.22 | 0.0003875 | 0.004122 | 5’ ccmK2 | - |
| slr1336 | 1.22 | 1.27E-08 | 5.07E-07 | SynCAX | TM |
| slr0770 | 1.22 | 0.0005199 | 0.005325 | slr0770 | - |
| sll0062 | 1.22 | 1.50E-06 | 3.59E-05 | sll0062 | - |
| slr1394 | 1.21 | 4.10E-09 | 1.81E-07 | slr1394 | - |
| slr0054 | 1.21 | 2.32E-07 | 7.08E-06 | dgkA | PM |
| sll1103 | 1.21 | 1.00E-07 | 3.34E-06 | gtrB | PM |
| slr2005 | 1.21 | 7.27E-07 | 1.89E-05 | slr2005 | - |
| slr0151 | 1.20 | 8.82E-07 | 2.22E-05 | slr0151 | PM |
| slr1508 | 1.12 | 4.11E-08 | 1.49E-06 | ktrE, dgdA | - |
| slr1275 | 1.19 | 5.60E-08 | 1.94E-06 | pilN | PM |
| sll1022 | 1.19 | 5.03E-08 | 1.77E-06 | sll1022 | - |
| sll1489 | 1.19 | 3.23E-07 | 9.37E-06 | cpmA | CP |
| slr0964 | 1.19 | 8.21E-08 | 2.79E-06 | slr0964 | PM |
| slr1150 | 1.18 | 0.000491 | 0.005092 | slr1150 | - |
| slr0363 | 1.18 | 0.000116 | 0.001536 | slr0363 | - |
| ssr1604 | 1.18 | 1.79E-06 | 4.20E-05 | rpL28 | - |
| slr0844 | 1.18 | 3.18E-10 | 1.79E-08 | ndhF1 | TM |
| sll0100 | 1.18 | 2.06E-07 | 6.33E-06 | ama | - |
| slr0772 | 1.17 | 1.25E-07 | 4.07E-06 | chlB | - |
| slr0351 | 1.17 | 2.91E-06 | 6.46E-05 | slr0351 | - |
| sll1712 | 1.16 | 8.05E-07 | 2.06E-05 | hu | - |
| slr0044 | 1.16 | 9.58E-07 | 2.38E-05 | cmpD | PM |
| sll1980 | 1.16 | 6.66E-07 | 1.76E-05 | trxA3 | TM |

| | | | | | |
|---|---|---|---|---|---|
| slr1984 | 1.16 | 1.19E-08 | 4.80E-07 | rps1b | - |
| smr0004 | 1.16 | 2.74E-06 | 6.16E-05 | psaI | TM |
| ssr3184 | 1.15 | 6.14E-06 | 0.000123 | fdx8 | - |
| slr2052 | 1.15 | 3.52E-07 | 1.01E-05 | slr2052 | CP |
| slr0145 | 1.15 | 0.000179 | 0.002172 | slr0145 | CP |
| slr0591 | 1.14 | 2.91E-07 | 8.63E-06 | nrdF | CP |
| slr1852 | 1.14 | 8.38E-06 | 0.000159 | slr1852 | CP |
| slr0043 | 1.14 | 8.03E-06 | 0.000154 | cmpC | PM |
| slr1853 | 1.14 | 0.000161 | 0.001999 | slr1853 | CP |
| slr1120 | 1.14 | 1.21E-06 | 2.96E-05 | pilD, hofD | PM |
| slr1145 | 1.14 | 1.54E-06 | 3.65E-05 | gltS | PM |
| slr1917 | 1.14 | 1.47E-08 | 5.69E-07 | slr1917 | - |
| slr1279 | 1.14 | 9.35E-11 | 6.14E-09 | ndhC | TM |
| sll1214 | 1.13 | 1.39E-05 | 0.000248 | cycI, ycf59, chlAI | TM |
| slr0250 | 1.12 | 2.89E-06 | 6.45E-05 | slr0250 | - |
| sll0185 | 1.12 | 2.63E-07 | 7.88E-06 | sll0185 | CP |
| sll1291 | 1.12 | 4.98E-06 | 0.000102 | rre12, taxP2 | PM |
| slr1438 | 1.12 | 1.01E-05 | 0.000188 | slr1438 | - |
| slr1176 | 1.11 | 5.61E-07 | 1.51E-05 | glgC, agp | CP |
| slr1055 | 1.11 | 1.19E-08 | 4.80E-07 | chlH | CP |
| sll0541 | 1.10 | 4.17E-07 | 1.16E-05 | desC, des9 | TM |
| sll1845 | 1.10 | 5.58E-07 | 1.51E-05 | sll1845 | - |
| sll1184 | 1.09 | 2.45E-06 | 5.60E-05 | ho1 | CP |
| ssr0390 | 1.09 | 8.66E-06 | 0.000164 | psaK1 | TM |
| ssr2087 | 1.09 | 0.000778 | 0.007347 | ssr2087 | - |
| sll1736 | 1.09 | 3.28E-05 | 0.000521 | sll1736 | - |
| slr1431 | 1.081 | 4.07E-07 | 1.14E-05 | slr1431 | PM |
| ssl1498 | 1.08 | 7.68E-05 | 0.001087 | ssl1498 | TM |
| slr1771 | 1.08 | 4.29E-07 | 1.18E-05 | slr1771 | CP |
| sll0686 | 1.08 | 5.22E-05 | 0.000771 | sll0686 | - |
| MYO_RS16815 | 1.08 | 0.000115 | 0.001524 | slr5056 | - |
| slr0342 | 1.08 | 1.49E-05 | 0.000265 | petB | TM |
| sll1219 | 1.08 | 1.37E-09 | 6.61E-08 | sll1219 | - |
| sll1098 | 1.07 | 3.22E-05 | 0.000514 | fus | CP |
| slr1138 | 1.07 | 5.76E-06 | 0.000116 | ctaEI | TM |
| sll0750 | 1.07 | 5.16E-06 | 0.000104 | hik8, sasA | - |
| sll0839 | 1.07 | 0.000725 | 0.007013 | sll0839 | PM |
| sll1049 | 1.07 | 1.72E-07 | 5.44E-06 | sll1049 | CP |
| sll1032 | 1.067 | 2.53E-05 | 0.000420 | ccmN | CBX |
| slr1547 | 1.06 | 7.81E-06 | 0.000151 | slr1547 | PM |
| sll0208 | 1.05 | 7.41E-08 | 2.53E-06 | fad | CP |
| slr1034 | 1.05 | 0.000303 | 0.003373 | ycf41 | TM |
| sll0222 | 1.04 | 4.64E-06 | 9.63E-05 | phoA | CP |
| sll1988 | 1.04 | 1.37E-05 | 0.000247 | hsp33 | CP |
| slr1838 | 1.04 | 1.38E-06 | 3.34E-05 | ccmK3 | CBX |

| | | | | | |
|---|---|---|---|---|---|
| sll0521 | 1.04 | 7.77E-06 | 0.000151 | ndhG | - |
| sll1688 | 1.04 | 2.82E-05 | 0.000459 | thrC | CP |
| slr1196 | 1.03 | 0.000129 | 0.001676 | slr1196 | ENV |
| sll1326 | 1.03 | 3.25E-06 | 7.08E-05 | atpA | TM |
| ssl0707 | 1.03 | 8.96E-07 | 2.24E-05 | glnB | - |
| slr2057 | 1.02 | 0.000355 | 0.00385 | apqZ | ENV |
| sll0169 | 1.02 | 1.41E-07 | 4.53E-06 | zipN, ftn2 | PM |
| sll1752 | 1.02 | 0.000133 | 0.001716 | sll1752 | TM |
| slr1280 | 1.02 | 2.23E-05 | 0.000379 | ndhK | TM |
| sll0681 | 1.02 | 5.75E-05 | 0.000839 | pstC | PM |
| sll0226 | 1.02 | 4.19E-06 | 8.80E-05 | ycf4 | TM |
| ssr3410 | 1.01 | 0.005997 | 0.036186 | mrpG | - |
| ssl2245 | 1.01 | 3.55E-05 | 0.000558 | ssl2245 | CP |
| ssr2799 | 1.01 | 8.14E-06 | 0.000156 | rpl27 | - |
| sll1832 | 1.01 | 1.11E-05 | 0.000206 | sll1832 | - |
| slr0058 | 1.01 | 7.77E-05 | 0.001094 | slr0058 | - |
| sll1899 | 1.01 | 2.06E-06 | 4.75E-05 | ctaB | - |
| ssl3093 | 1.01 | 7.09E-05 | 0.001012 | cpcD | TM |
| sll1031 | 1.01 | 5.06E-06 | 0.000103 | ccmM | CBX |
| sll0891 | 1.01 | 4.97E-06 | 0.000102 | citH, mdh | CP |
| slr2136 | 1.00 | 1.40E-05 | 0.000251 | gcpE | CP |
| slr0346 | 1.00 | 0.000164 | 0.002018 | rnc | CP |
| slr1906 | 1.00 | 8.63E-05 | 0.001198 | slr1906 | - |

**Table S2. Proteins significantly enriched in the co-IP with Rbp3 after identification by MS.** The protein and gene names are given, together with the enrichment ($log_2FC$), and the adjusted *p* value compared to the sfGFP negative control, followed by the respective uniprot ID. The underlying data are average LFQ counts from three biological replicates each for the co-IP with Rbp3 and sfGFP, respectively. For the complete list of detected proteins, see **Table S9.** This table extends the data in **Fig. 4A**.

| Protein | Gene | $log_2FC$ | adj. p-value | Uniprot ID |
|---|---|---|---|---|
| Slr1859 | slr1859 | Only in Rbp3 IP | Only in Rbp3 IP | P73609 |
| Slr1870 | slr1870 | Only in Rbp3 IP | Only in Rbp3 IP | P73622 |
| Rbp3 | slr0193 | 14.59 | 1.05E-02 | P73124 |
| Slr0650 | slr0650 | 3.97 | 1.15E-02 | P74485 |
| Dfa2 | dfa2 | 3.81 | 1.84E-02 | Q55765 |
| Rplb | rplB | 3.72 | 1.60E-02 | Q55730 |
| Sll0518 | sll0518 | 3.69 | 1.20E-02 | P72723 |
| Rplm | rplM | 3.68 | 1.49E-02 | P73317 |
| Rplw | rplW | 3.54 | 5.94E-03 | Q55466 |
| Rpso | rpsO | 3.49 | 1.60E-02 | P73294 |
| Rpls | rplS | 3.46 | 2.79E-02 | P73318 |
| Rpsh | rpsH | 3.39 | 1.90E-02 | P72866 |
| Rplq | rplQ | 3.38 | 1.05E-02 | P36239 |
| Rpln | rplN | 3.36 | 2.24E-02 | P73307 |
| Rpsp | rpsP | 3.34 | 1.60E-02 | P73296 |
| Rpsd | rpsD | 3.33 | 2.70E-02 | P73310 |
| Rplo | rplO | 3.31 | 4.02E-02 | P74410 |
| Rpmc | rpmC | 3.29 | 1.49E-02 | P48939 |
| Rpli | rplI | 3.28 | 1.48E-02 | P73303 |
| Rple | rplE | 3.24 | 1.20E-02 | P73312 |
| Rpse | rpsE | 3.23 | 2.31E-02 | P42352 |
| Rpsi | rpsI | 3.22 | 1.20E-02 | P73308 |
| Pnp | pnp | 3.18 | 1.08E-02 | P73304 |
| Rpsk | rpsK | 3.17 | 1.60E-02 | P73293 |
| Rps1A | rps1A | 3.17 | 5.94E-03 | P72659 |
| Rplc | rplC | 3.17 | 1.48E-02 | P73298 |
| Rpsm | rpsM | 3.11 | 1.49E-02 | P73530 |
| Rpma | rpmA | 3.09 | 3.77E-02 | P73320 |
| Rpsc | rpsC | 3.08 | 1.60E-02 | P73299 |
| Rpld | rplD | 3.05 | 4.73E-02 | P74267 |
| Rplp | rplP | 3.00 | 1.53E-02 | P73314 |
| Rpst | rpsT | 2.97 | 1.90E-02 | P73319 |
| Rpla | rplA | 2.97 | 1.84E-02 | P73313 |
| Rpsg | rpsG | 2.93 | 1.60E-02 | P73336 |

| | | | | |
|---|---|---|---|---|
| Rplv | rplV | 2.91 | 1.60E-02 | P36236 |
| Rpsf | rpsF | 2.87 | 3.33E-03 | P74229 |
| Rpsb | rpsB | 2.80 | 1.60E-02 | P73315 |
| Rplf | rplF | 2.78 | 2.97E-02 | P73636 |
| Dfa4 | dfa4 | 2.70 | 1.60E-02 | P74071 |
| Rpss | rpsS | 2.59 | 3.75E-02 | P73306 |
| Rpmb | rpmB | 2.58 | 1.60E-02 | P72721 |
| Rplu | rplU | 2.52 | 2.24E-02 | P73316 |
| Rpsr | rpsR | 2.51 | 2.79E-02 | P72851 |
| Rpsj | rpsJ | 2.43 | 2.24E-02 | P74266 |
| Rpme | rpmE | 2.37 | 3.75E-02 | P48946 |
| Rpsq | rpsQ | 2.31 | 3.75E-02 | P74226 |
| Rplr | rplR | 2.23 | 2.90E-02 | P73292 |
| Rpsu | rpsU | 2.04 | 2.97E-02 | P73311 |
| Rplj | rplJ | 2.04 | 1.60E-02 | P73305 |
| Slr0635 | slr0635 | 2.01 | 5.94E-03 | P48949 |
| Slr1034 | slr1034 | 1.96 | 1.60E-02 | P23350 |
| Gspe | gspE | 1.93 | 1.92E-02 | Q55712 |
| Ssb | ssb | 1.90 | 5.94E-03 | P73145 |
| Slr1854 | slr1854 | 1.85 | 4.48E-02 | Q55155 |
| Rnj | rnj | 1.59 | 3.59E-02 | Q55499 |
| Nira | nirA | 1.57 | 4.35E-02 | P73605 |
| Slr0322 | slr0322 | 1.41 | 2.24E-02 | P54123 |
| Rpll | rplL | 1.36 | 3.75E-02 | Q55366 |
| Slr1223 | slr1223 | 1.24 | 5.94E-03 | Q55527 |
| Dnak2 | dnaK2 | 1.24 | 1.60E-02 | P23349 |
| Prs | prs | 1.21 | 4.88E-02 | P73467 |
| Acpp | acpP | 1.18 | 4.40E-02 | P22358 |
| Cpcl | cpcL | 1.13 | 4.42E-02 | Q55848 |
| Slr0006 | slr0006 | 1.09 | 4.35E-02 | P20804 |
| Carb | carB | 1.01 | 3.36E-02 | P74625 |

**Table S3. Transcriptomic differences in the comparison of *Δrpb2Δrpb3* versus wild type under non-stress conditions and after shift to 20°C.** This table extends the data in **Fig. 5C, D**.

See separate Excel spreadsheet.

**Table S4. Transcriptomic differences in the comparison of *Δrpb2Δrpb3* versus wild type under non-stress conditions and after shift to high light.** This table extends the data in **Fig. 5E, F**.

See separate Excel spreadsheet.

**Table S5. Primers and plasmids used in this work.** The oligonucleotide probes used for mRNA-FISH are listed in a separate Excel spreadsheet (**Table S6**).

| **Oligonucleotide primers** | | |
|---|---|---|
| **Name** | **Sequence** | **Description** |
| sfGFP_fwd | GCTCTACAAAATGGATTATAAAGATCATGATGG | Amplification of *sfGFP* from pXG10-SF with overlaps for AQUA cloning into pJet1.2-pPetE |
| sfGFP_rev | CCCGGCGGCAACCGAGCGAACTATTTATCATCATCATCTTTATAATCAATATC | |
| pJet1.2_PpetE_fwd | TTCGCTCGGTTGCCGCCG | Linearization of pJet1.2-PpetE-*rbp3* excluding *rbp3* |
| pJet1.2_PpetE_rev | ACTTCTTGGCGATTGTATCTATAGGGACTTGTG | |
| pJET1.2_3xFLAG_in_tooP_fwd | TCATGATATTGATTATAAAGATGATGATGATAAATAGTTCGCTCGGTTGCCGCCG | Introduction of 3xFLAG tag at the GFP C-terminus sequence by inverse PCR |
| pJet1.2_sfGFP_3xFLAG_in_rev | TCTTTATAATCGCCATCATGATCTTTATAATCTTTGTAGAGCTCATCCATGCCATGTG | |
| AQUA_sfGFP_fwd | TCCTGCAGGTCGACTCTAGACTGGGCCTACTGGGCTATTC | Primer for pJet1.2::PpetE::*sfGFP::* 3xFLAG with overlaps to pVZ322 digested with XbaI/HindIII |
| AQUA_sfGFP_rev | GGAAAGAAATGCATAAGCTTAATAAAAAACGCCCGGCG | |
| *HindIII*-Slr0193_fwd | AAGCTTatgcttattcccgtttgattg | Amplification of *rbp3* from the genome, introducing digestion sites for *HindIII* and *XhoI* |
| *XhoI*-Slr0193_rev | CTCGAGgttttttattaaactctaaacaggacaaag | |
| psaA_probe_fwd | TCGACCGGACTTTAGCTAG | Primer for nucleotide probe for *psaA* |
| psaA_T7_probe_rev | TAATACGACTCACTATAGGGCAAAGACCACAGCGAGGTG | |
| pGEX-6P-1_inv-f | AATGGTTTCTTAGACGTCAGGTGGC | |
| pGEX-6P-1_inv-r | GAATTCCGGGGATCCCAGGG | |
| Rbp3_iVEC-f | CTGGGATCCCCGGAATTCATGTCCATTCGTCTCTACGTCG | |
| Rbp3_iVEC-r | CTGACGTCTAAGAAACCATTCTACTGGGCCGCTGTCAGTTTTT | |

| | | |
|---|---|---|
| FP | ACGATGGGGAGAAAGAAACCGTAG | |
| RP | GACCAATCAGAGTGACATTGGGCA | |
| Seg_rbp3_fwd | ACGATGGGGAGAAAGAAACCGTAG | Colony PCR delta rbp3 |
| Seg_rbp3_rev | GACCAATCAGAGTGACATTGGGCA | |
| 51_Rbp3_Linker_sfGFP_fw | CTGACAGCGGCCCAGGGATCCGCTGGCTCCG | Amplification of *sfGFP* sequence |
| Rbp3_3'UTR_GFP _rev | GGAAAACCTGTGGACTTATTTGTAGAGCTCATCC | |
| Rbp3_3'UTR_GFP _fw | GGATGAGCTCTACAAATAAGTCCACAGGTTTTCC | Amplification of pVZ322: pRbp3-*rbp3* |
| 50_Rbp3_Linker_sfGFP_rev | CGGAGCCAGCGGATCCCTGGGCCGCTGTCAG | |
| 56_rbp-sfgfp-pVZ_cPCR_fw | TGGACGGTAACCGAGTTCGCGG | Successful integration of *sfGFP* in pVZ322::*rbp3* plasmid |
| 57_rbp-sfgfp-pVZ_cPCR_rev | GCCCTGCTGCGTAACATCGTTG | |
| **Plasmids** | | |
| **Name** | **Reference** | |
| pJet1.2 | Thermo Fisher Scientific "CloneJET" | |
| pVZ322 | (12) | |
| pXG10-SF | (13) | |
| pVZ322::PpetE::rbp3::3xFLAG | (14) | |
| pVZ322::PpetE::sfGFP::3xFLAG | This study | |

**Table S6. Oligonucleotide probes used for mRNA-FISH.**

See separate Excel spreadsheet.

**Table S7. Details of the statistical analysis of colony sizes in Fig. 1A, B.**

| WT vs mutant<br>t-Test: Two-Sample Assuming Equal Variances | | |
|---|---|---|
| | *Variable 1* | *Variable 2* |
| Mean | 0.62725676 | 0.38187826 |
| Variance | 0.01104565 | 0.01050913 |
| Observations | 74 | 115 |
| Pooled Variance | 0.01071857 | |
| Hypothesized Mean Difference | 0 | |
| df | 187 | |
| t Stat | 15.9038391 | |
| P(T<=t) one-tail | 6.9751E-37 | |
| t Critical one-tail | 1.65304289 | |
| P(T<=t) two-tail | 1.395E-36 | |
| t Critical two-tail | 1.97273103 | |

| Comp. Vs. Mutant<br>t-Test: Two-Sample Assuming Equal Variances | | |
|---|---|---|
| | *Variable 1* | *Variable 2* |
| Mean | 0.61985507 | 0.38187826 |
| Variance | 0.01056168 | 0.01050913 |
| Observations | 69 | 115 |
| Pooled Variance | 0.01052876 | |
| Hypothesized Mean Difference | 0 | |
| df | 182 | |
| t Stat | 15.2303724 | |
| P(T<=t) one-tail | 1.3095E-34 | |
| t Critical one-tail | 1.65326902 | |
| P(T<=t) two-tail | 2.6189E-34 | |
| t Critical two-tail | 1.97308408 | |

**Table S8. Details of the statistical analysis for the quantification of *psaA* mRNA fluorescence intensity comparing wild type, Δ*rbp3* and Δ*rbp3* + Rbp3-sfGFP strains.** The fluorescence intensities were determined in background corrected images. This Table extends **Fig. 2G.**

| WT vs Δrbp3 | | |
|---|---|---|
| t-Test: Two-Sample Assuming Equal Variances | | |
| | *Variable 1* | *Variable 2* |
| Mean | 166.8031 | 76.71078 |
| Variance | 5175.055 | 1050.244 |
| Observations | 232 | 231 |
| Pooled Variance | 3117.123 | |
| Hypothesized Mean Difference | 0 | |
| df | 461 | |
| t Stat | 17.36083 | |
| P(T<=t) one-tail | 1.29E-52 | |
| t Critical one-tail | 1.648166 | |
| P(T<=t) two-tail | 2.57E-52 | |
| t Critical two-tail | 1.965123 | |

| RBP3-sfGFP vs. Δrbp3 | | |
|---|---|---|
| t-Test: Two-Sample Assuming Equal Variances | | |
| | *Variable 1* | *Variable 2* |
| Mean | 206.0466 | 76.71078 |
| Variance | 7227.565 | 1050.244 |
| Observations | 232 | 231 |
| Pooled Variance | 4145.604 | |
| Hypothesized Mean Difference | 0 | |
| df | 461 | |
| t Stat | 21.61148 | |
| P(T<=t) one-tail | 2.37E-72 | |
| t Critical one-tail | 1.648166 | |
| P(T<=t) two-tail | 4.73E-72 | |
| t Critical two-tail | 1.965123 | |

**Table S9**. **Complete list of proteins detected by mass spectrometry in the co-IP experiments with Rbp3.** This table extends the data in **Fig. 4A, B** and in **Table S2**. See separate Excel spreadsheet.

**Table S10**. Details of data analysis - hierarchical clustering in **Fig. 4B** (Perseus v2.0.11 (8)).

| | | |
|---|---|---|
| **Presets** | $log_2$ transformed LFQ intensities | |
| | ribosomal proteins were selected for clustering | |
| | No imputation of missing values | |
| **Rows** | | |
| Distance | Euclidean | |
| Linkage | Average | |
| Constraint | None | |
| Preprocess with k-means | Yes | |
| | Number of clusters | 300 |
| | Maximal number of Iterations | 10 |
| | Number of restarts | 1 |
| **Columns** | | |
| Distance | Euclidean | |
| Linkage | Average | |
| Constraint | None | |
| Preprocess with k-means | Yes | |
| | Number of clusters | 300 |
| | Maximal number of Iterations | 10 |
| | Number of restarts | 1 |

**Table S11. Calculation of Pearson's correlation coefficients in the colocalization of fluorescence signals (numeric values).** This Table extends **Fig. 4C, D.**
See separate Excel spreadsheet.

**Table S12. Details of the statistical analysis of cell sizes in Fig. S3.**

| WT vs mutant<br>t-Test: Two-Sample Assuming Equal Variances | | |
|---|---|---|
| | *Variable 1* | *Variable 2* |
| Mean | 2.806491 | 2.215185 |
| Variance | 0.508385 | 0.258217 |
| Observations | 322 | 432 |
| Pooled Variance | 0.365004 | |
| Hypothesized Mean Difference | 0 | |
| df | 752 | |
| t Stat | 13.29373 | |
| P(T<=t) one-tail | 1.13E-36 | |
| t Critical one-tail | 1.646882 | |
| P(T<=t) two-tail | 2.26E-36 | |
| t Critical two-tail | 1.963124 | |

| Comp. vs mutant<br>t-Test: Two-Sample Assuming Equal Variances | | |
|---|---|---|
| | *Variable 1* | *Variable 2* |
| Mean | 3.056259 | 2.215185 |
| Variance | 0.433245 | 0.258217 |
| Observations | 286 | 432 |
| Pooled Variance | 0.327886 | |
| Hypothesized Mean Difference | 0 | |
| df | 716 | |
| t Stat | 19.26793 | |
| P(T<=t) one-tail | 2.87E-67 | |
| t Critical one-tail | 1.646985 | |
| P(T<=t) two-tail | 5.74E-67 | |
| t Critical two-tail | 1.963283 | |

### Supplementary Figures

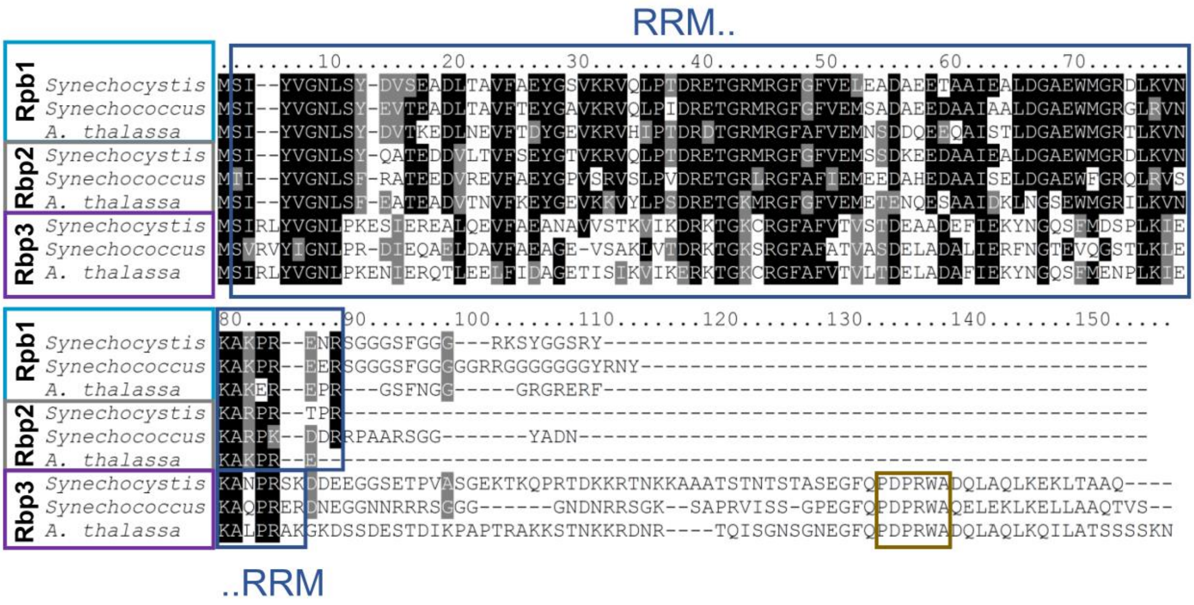

**Figure S1. Sequence comparison of RRM domain proteins in cyanobacteria.** Three main families can be distinguished, Rbp1, Rbp2 and Rbp3. These proteins are encoded by small gene families in all cyanobacteria, including derivatives such as the most genome-streamlined $N_2$-fixing nitroplast (15), with a genome size of just 1.44 Mbp (16). The multiple sequence alignment shows the three respective RRM domain-containing RBPs each from *Synechocystis* 6803, *Synechococcus elongatus* sp. PCC 7942 and the nitroplast (*Candidatus* Atelocyanobacterium thalassa strain ALOHA). The RRM domain and PDPRWA motif are boxed. The Rbp1 family is characterized by a glycine-rich C-terminal domain. Members of the Rbp2 family represent the shortest RRM-domain proteins in cyanobacteria, encompassing just a single RRM domain, while the Rbp3 family proteins are longer and feature a conserved "PDPRWA" motif near the C-terminus (17, 18).

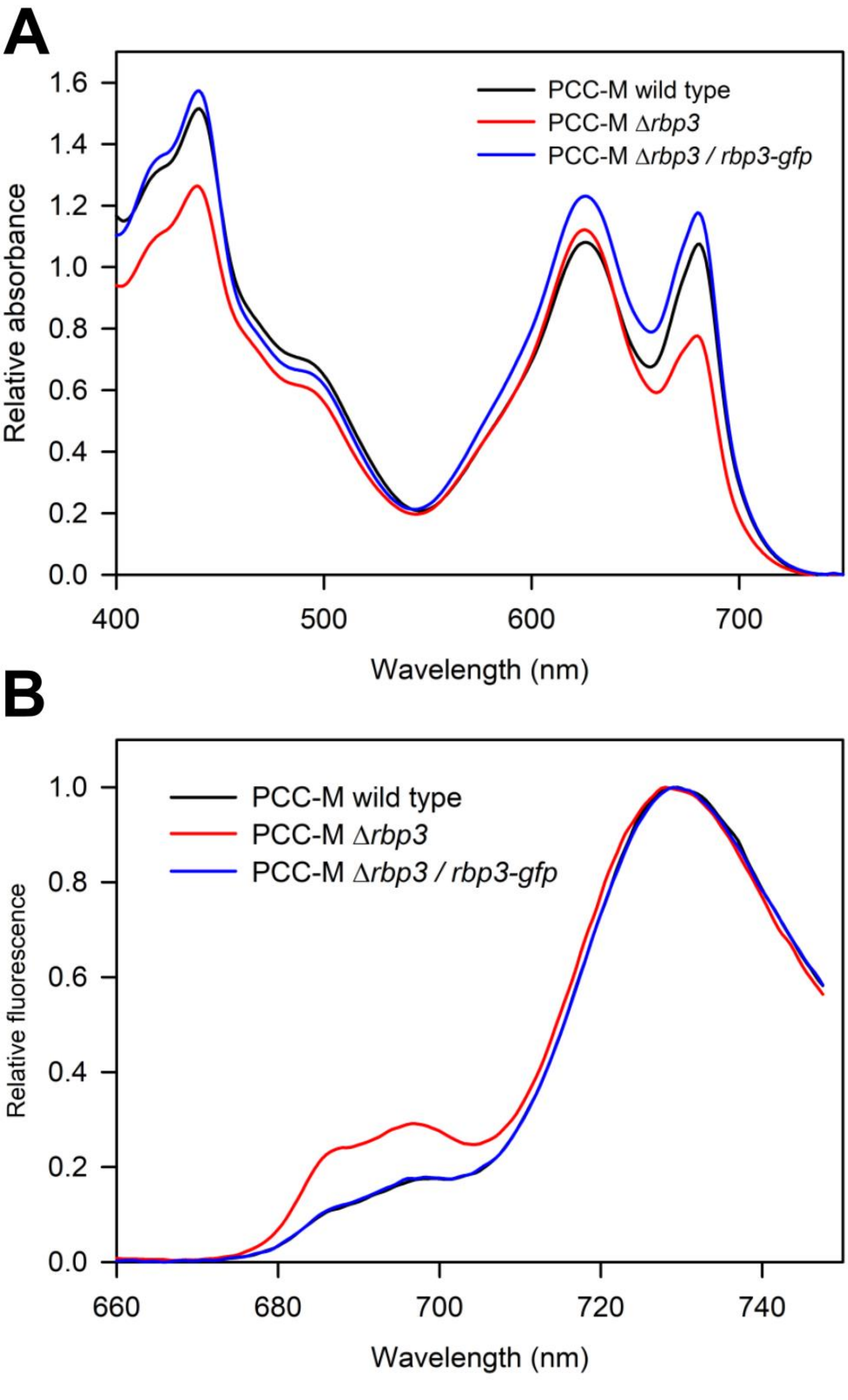

**Figure S2. Spectral characterization of *Synechocystis* 6803 PCC-M wild type, Δ*rbp3* mutant and a complementation strain expressing an Rbp3-sfGFP fusion protein. (A)** Room-temperature absorption spectra of cell suspensions for the PCC-M wild type, Δ*rbp3* and Δ*rbp3*+*rbp3*-sfGFP, normalized at 750 nm. **(B)** 77K fluorescence emission spectra with excitation at 435 nm and normalized to the PSI peak for the same strains as in panel A. This figure is an extension of **Fig. 2**.

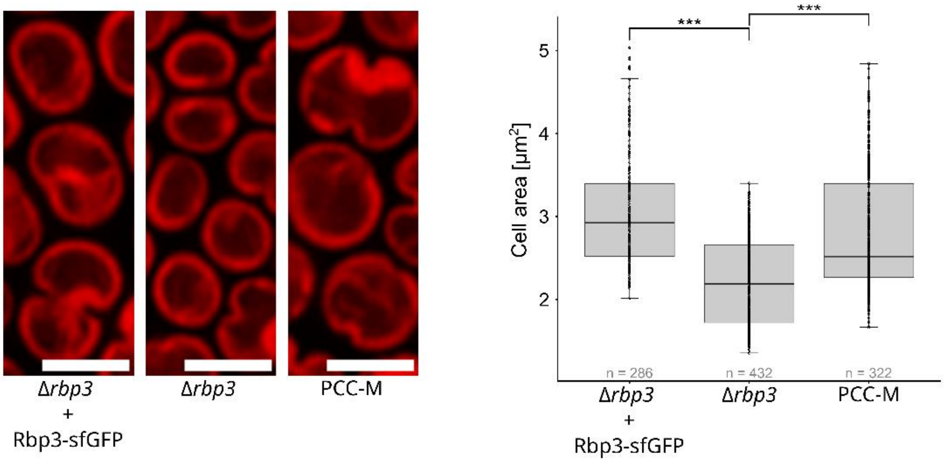

**Figure S3. Confocal images of *Synechocystis* PCC-M wild type, *Δrbp3* and *Δrbp3*+Rbp3-sfGFP cells showing thylakoid membrane chlorophyll fluorescence (red).** The cell size was calculated and compared between the strains (right panel). n = number of cells; *** significant difference between the strains ($p < 0.001$). Significances were calculated by the students t-test assuming equal variances, for the numeric values see **Table S12.** This figure extends **Fig. 2F**.

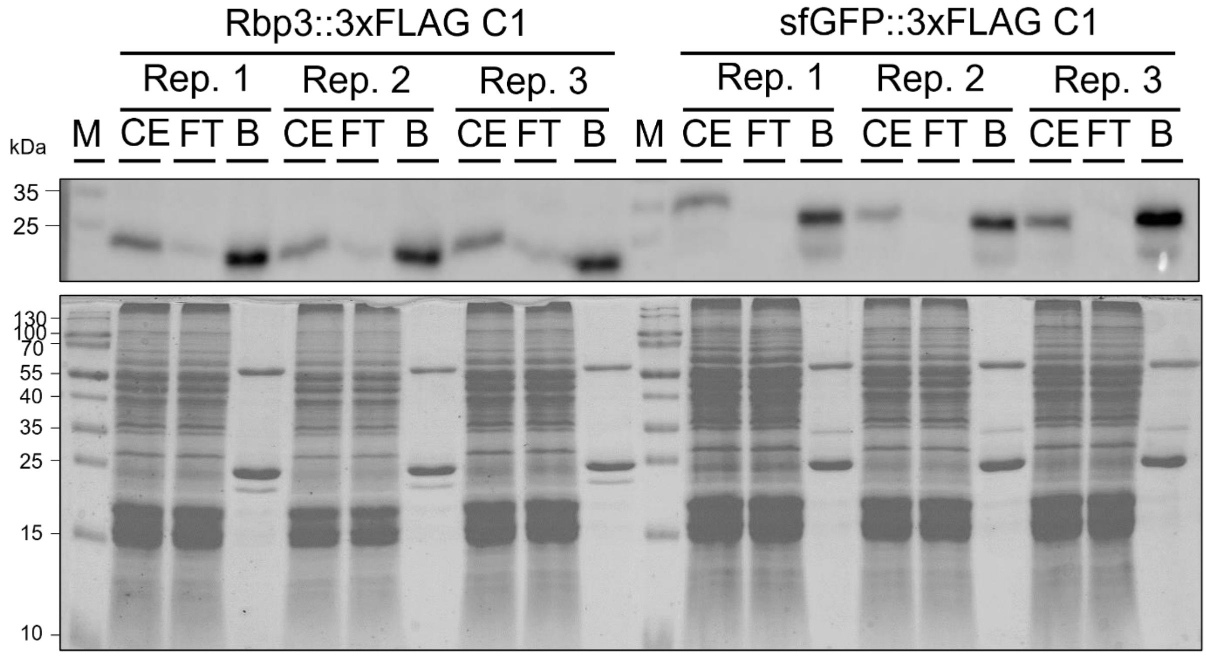

**Figure S4. Western blot (upper panel) and coomassie-stained SDS-PAGE (lower panel) of the cross-linked immunoprecipitation of strains containing a pVZ322 plasmid harboring following constructs: $P_{petE}$_rbp3_3×FLAG and $P_{petE}$_sfGFP_3×FLAG from which upon induction either Rbp3_3×FLAG or sfGFP_3×FLAG was expressed.** The experiment was performed in technical triplicates (Rep. 1-3). Purification of the 3×FLAG-tagged proteins was done using anti FLAG M2 magnetic beads (Sigma Aldrich). Proteins were separated on a 15% SDS-PAA gel. Ten µL of bead suspension after washing were mixed 1:1 with 2x protein loading buffer. After heat elution the supernatant was loaded in the bead fraction. For all other fractions 10 µL of either 1:1 diluted crude extract or 1:1 diluted flow through was loaded. Bands visible at ~20 kDa belong to Rbp3-3xFLAG, bands at ~35 kDa to sfGFP-3xFLAG. Bands present in all bead fractions at ~25 and ~55 kDa belong to the chains of the anti-FLAG antibody that was denatured during heat elution of beads. Marker (M): PageRuler (Thermo Fischer Scientific), CE: crude extract, FT: flow through, B: bead fraction. In the upper panel, the ANTI-FLAG M2-peroxidase antiserum (Sigma Aldrich) was used at a titer of 1:5,000.

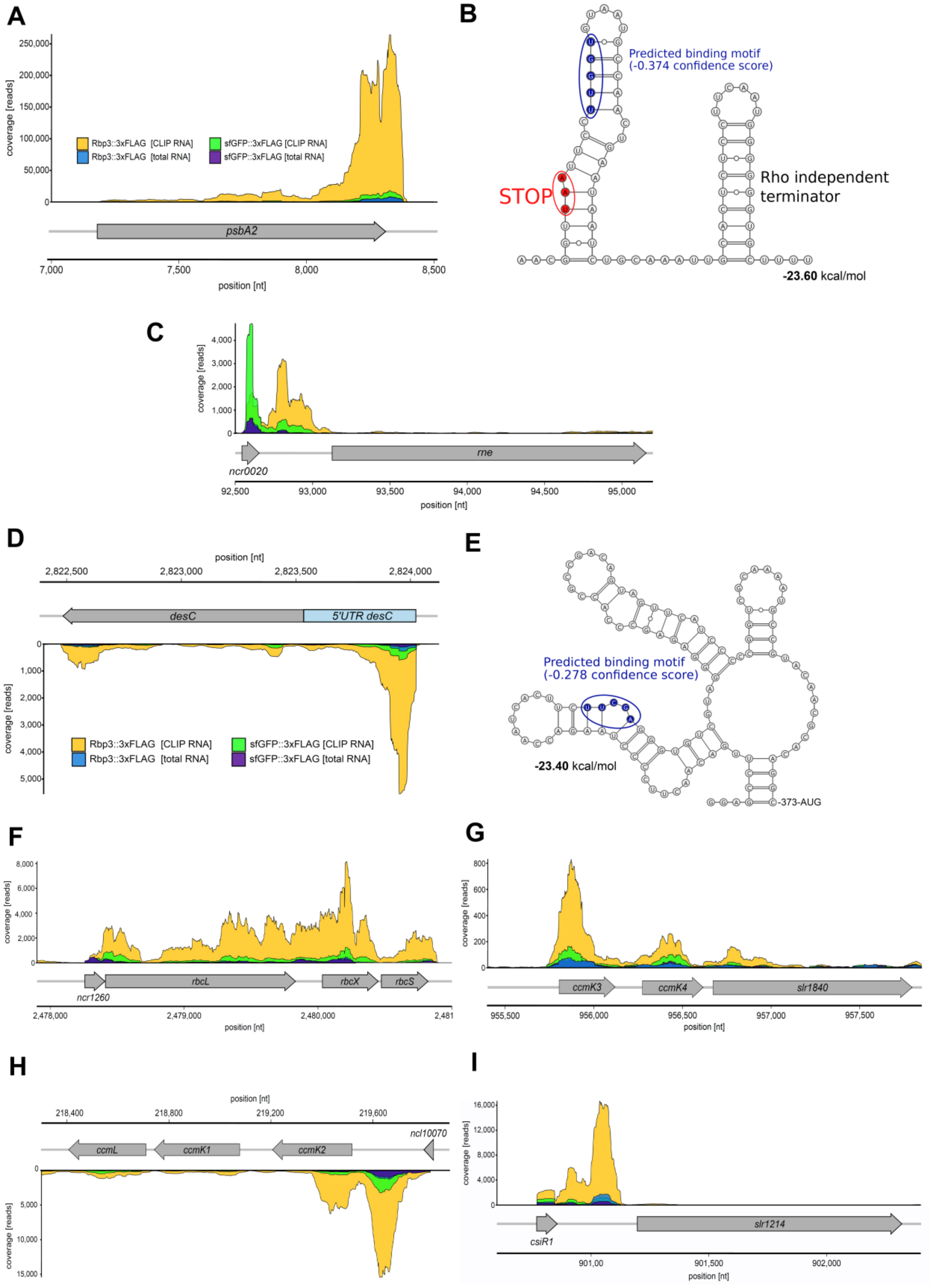

**Figure S5. Coverage of enriched mRNA reveals peaks in distinct areas of the transcripts. (A)** Read coverage alongside the *psbA2* gene and **(B)** secondary structure prediction for the peak area. Note, that we chose the *psbA2* sequence for visualization, but the two genes *psbA2* and *psbA3* are nearly identical and were not differentiated based on mapped cDNA reads. **(C)** The *rne* gene encoding RNase E. **(D)** The *desC* (*sll0541*) gene encoding delta 9 acyl-lipid desaturase. **(E)** Secondary structure prediction for the peak area in the *desC* 5'UTR. Putative RRM binding sites predicted by the RRMScorer algorithm are circled in blue (19). **(F)** The *rbcLXS* tricistron, encoding the large and small subunits of RuBisCO and the RbcX chaperone. **(G)** The *ccmK3K4* locus. **(H)** The *ccmK1K2* locus. **(I)** The *csiR1-slr1214* locus. For other explanations, see **Fig. 3** in the main document.
